# EWSR1::ATF1 hyperactivation via TRIM11 depletion triggers lethal metabolic rewiring in clear cell sarcoma

**DOI:** 10.64898/2026.11.06.764260

**Authors:** Wenxin Chao, Kilian Bales, Kamini Kaushal, Laura Mayerhofer, Julien Picotto, Akanksha Farswan, Christopher Davies, Dawid Krzeciesa, Susanne Grunewald, Francisco Yanqui-Rivera, Nicola Biondi, Liang Chen, Adarsh Raveendran, Anushka Deshpande, Kwang Seok Lee, Martin Schneider, Dario Frey, Shiva Ahmadi, Mohamad Saoud, Michael Buettner, Karsten Richter, Michelle Neßling, Claudia Scholl, Stefan Fröhling, Ashraf Yusuf Rangrez, Kevin Jones, Sumathi Vaiyapuri, Daniel Hughes, Sandra Strauss, Gunhild Mechtersheimer, Christiane A. Opitz, Nischalan Pillay, Mikolaj Slabicki, Sebastian Bauer, Priya Chudasama

## Abstract

Clear cell sarcoma (CCS) is an ultra-rare and aggressive soft tissue sarcoma driven by the EWSR1::ATF1 fusion oncoprotein. Currently, no effective and well-tolerated treatment options are available to CCS patients. Direct inhibition of fusion transcription factors is precluded by the absence of defined ligand-binding pockets, whereas their abundance is under the control of the ubiquitin-proteasome system (UPS), offering an alternative avenue for biology-guided treatment. Here we show the abundance of EWSR1::ATF1 is tightly regulated by the UPS, and that release of this constraint is lethal to CCS cells. A reporter-based CRISPR/Cas9 knockout screen of UPS components identified TRIM11 as an E3 ligase that regulates turnover of EWSR1::ATF1, which it recognises as a neosubstrate without altering wild-type EWSR1 or ATF1. Depletion of TRIM11 increased nuclear fusion protein abundance, amplified the fusion-driven transcriptional programs and reduced viability. Proteomic analysis identified the cataplerotic enzyme phosphoenolpyruvate carboxykinase 1 (PCK1), a direct EWSR1::ATF1 target, as the key mediator of lethal effects of fusion hyperactivation. Extracellular flux analysis, isotope tracing and quantitative metabolomics revealed excessive cataplerotic carbon withdrawal and impaired oxidative phosphorylation as the underlying mechanism. In patient tumors, high PCK1 expression associated with metastatic transcriptome programs and disease presentation, as well as shorter survival. Taken together, we report that hyperactivation of EWSR1::ATF1 opens a metabolic vulnerability, establishing TRIM11 as an actionable dependency and PCK1 as a candidate prognostic biomarker in CCS.

## Introduction

Clear cell sarcoma (CCS) is a rare, aggressive soft tissue sarcoma that predominantly affects adolescents and young adults, and carries a poor prognosis^1,2^. It often arises in the deep soft tissues of the distal extremities and metastasizes frequently to lymph nodes and lung. Surgical resection remains the only treatment with curative potential, while cytotoxic chemotherapy is considered ineffective. For patients with metastatic disease, currently clinical trials are considered standard of care. In-field relapses are more frequently observed following radiotherapy and the role of perioperative radiotherapy remains uncertain^3,4^. Although historically termed malignant melanoma of soft parts on the basis of shared melanocytic differentiation markers, CCS is distinguished molecularly by the recurrent t(12;22)(q13;q12) translocation, which generates EWSR1::ATF1, the fusion transcription factor that acts as the oncogenic driver of the disease^5,6^. CCS further belongs to the group of ultra-rare sarcomas, defined by an annual incidence at or below one case per million, a rarity that restricts accrual to histotype-specific trials, limits the accumulation of biological knowledge, and impedes authorization of new agents^7^. Accordingly, CCS remains a challenging entity with urgent unmet medical need characterized by aggressive clinical course and limited therapeutic options.

Current understanding of CCS biology centers largely on the transcriptional and epigenetic functions of the oncogenic fusion protein. Recent studies have shown that EWSR1::ATF1 reprograms chromatin accessibility, establishes tumor-specific enhancer networks and reshapes higher-order chromatin architecture to sustain the oncogenic program^8–10^. Therapeutic strategies have focused on downstream effectors of EWSR1::ATF1, including tyrosine kinase inhibitors acting on MET- and VEGFR-associated signaling, which have yielded limited clinical benefit^11,12^. Directly targeting EWSR1::ATF1 therefore represents an attractive strategy to overcome the limitations of targeting secondary oncogenic pathways in CCS. Unlike kinase-domain-containing fusion oncoproteins such as BCR::ABL1, ETV6::NTRK3 and EML4::ALK, which drive tumorigenesis through constitutive activation of retained tyrosine kinase domains^13–15^, EWSR1::ATF1 functions primarily as an oncogenic transcription factor. Its lack of conventional ligand-binding pockets has rendered it, like many oncogenic transcription factors, largely inaccessible to traditional small-molecule inhibition.

Cancer cells depend on tightly regulated protein turnover, creating opportunities to therapeutically manipulate the ubiquitin-proteasome system (UPS). Targeted degradation strategies, including “molecular glues” and proteolysis targeting chimeras (PROTACs), exploit E3 ubiquitin ligases to eliminate otherwise “undruggable” oncoproteins (PMID: 19029980; PMID: 24292625; PMID: 31901786; PMID: 11438690). Conversely, disrupting oncoprotein turnover can drive its abundance beyond a tolerable threshold, resulting in excessive oncogenic signaling and oncogene-induced senescence (PMID: 30475204; PMID: 34329586). Thus, manipulating protein turnover offers two complementary strategies: degrading an oncogenic driver or hyperactivating it beyond the level tolerated by cancer cells.

The ubiquitin-proteasome system (UPS) is a key intracellular protein degradation and turnover system (PMID: 18930136). Cancer cells depend on this control, and perturbation of UPS-mediated turnover has emerged as a strategy for tackling “undruggable” proteins of interest. “Molecular glues” and proteolysis targeting chimeras (PROTACs) achieve this by recruiting the oncogenic protein to an E3 ligase for degradation (PMID: 19029980, PMID: 24292625, PMID: 31901786, PMID: 11438690), whereas disrupting the turnover that constrains its abundance acts in an opposite direction by driving the oncogenic program past the level cells tolerate, and could generate a vulnerability specific to hyperactivation (PMID: 30475204, PMID: 34329586).

In this study, we aimed to uncover the mechanisms governing the stability of EWSR1::ATF1, and identified therapeutic entry points within protein homeostasis pathways. We identified Tripartite Motif Containing 11 (TRIM11), an E3 ligase, as a regulator of fusion protein abundance. We show that depletion of TRIM11 increased fusion protein abundance and elicited a metabolic shift accompanied by mitochondrial dysfunction and loss of cell viability. Mechanistically, we connected this vulnerability to excessive induction of PCK1, and the cataplerotic efflux of tricarboxylic acid cycle intermediates, resulting in altered cellular energy metabolism and reduced CCS cell growth. Clinically, higher PCK1 expression levels correlated with metastatic features and overall survival in CCS, extending its role as a candidate prognostic biomarker.

## Results

### TRIM11 regulates the turnover of EWSR1::ATF1

To determine whether EWSR1::ATF1 protein abundance is regulated by the UPS, we used the p-Artichoke reporter system^16,17^ in HEK293T Cas9 cells. This system expresses eGFP-tagged fusion proteins alongside mCherry as an internal expression control, allowing relative protein stability to be quantified by the eGFP/mCherry ratio (Figure 1A). The fusion protein driving Ewing sarcoma, namely EWSR1::FLI1, whose turnover is known to depend on the UPS^18^, was included as a positive control, while FLAG served as a negative control. Treatment with either the proteasome inhibitor MG132 or the E1-activating enzyme inhibitor MLN7243 rescued the degradation of both EWSR1::ATF1 and EWSR1::FLI1, whereas the FLAG reporter remained unchanged (Figure 1B). These findings demonstrate that EWSR1::ATF1 abundance is constrained by the UPS-mediated turnover.

**Figure 1.**
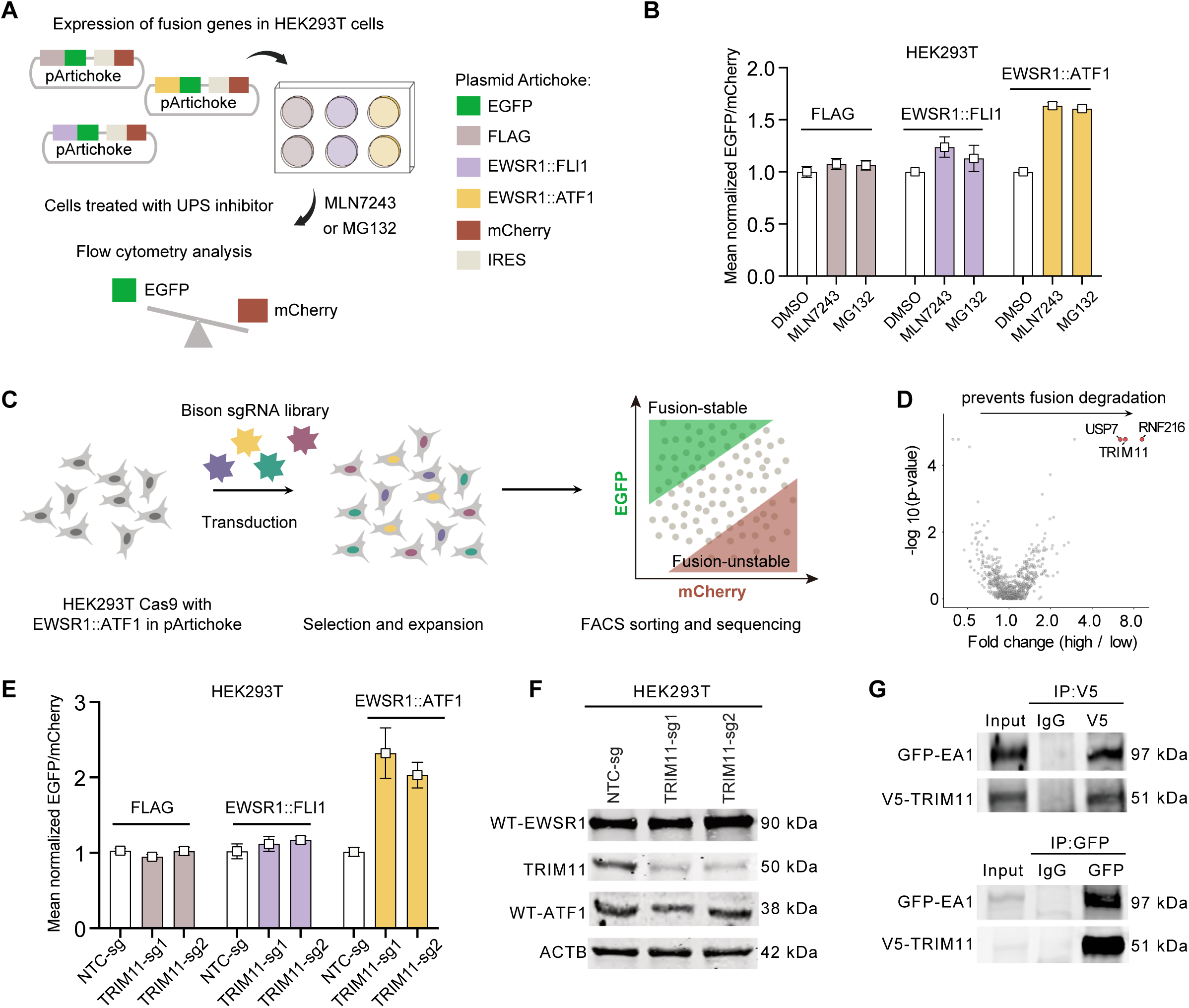
Ubiquitin–proteasome system (UPS) focused CRISPR screen identified TRIM11 as regulator of EWSR1::ATF1 fusion stability. (A) Schematic illustrating the design of the reporter plasmid pArtichoke and the workflow used to assess UPS-dependent fusion protein degradation. EGFP, enhanced green fluorescent protein; IRES, internal ribosome entry site. (B) Flow cytometric analysis of EWSR1::ATF1 reporter enrichment following treatment with the proteasome inhibitor MG132 (0.5 µM) or the E1 inhibitor MLN7243 (1 µM) for 2 h, relative to the dimethyl sulfoxide (DMSO) control. FLAG and EWSR1::FLI1 served as negative and positive controls. n = 3 biological replicates. (C) Schematic of the reporter-based Bison CRISPR/Cas9 knockout screen workflow. sgRNA, single-guide RNA; FACS, fluorescence-activated cell sorting. (D) Volcano plot showing gene-level enrichment in the EWSR1::ATF1 reporter screen comparing high versus low fluorescence gates. The x-axis shows the median fold change of normalized sgRNA counts in the high gate relative to the low gate. The y-axis shows the -log10 empirical P value from rank-based permutation testing. Each point represents one gene. TRIM11 is highlighted in red. (E) Flow cytometric analysis of fusion enrichment upon TRIM11 knockout (TKO) compared with non-targeting control (NTC) in HEK293T with EWSR1::ATF1 pArtichoke. n = 3 biological replicates. (F) Western blot analysis of wild-type EWSR1 (WT-EWSR1) and wild-type ATF1 (WT-ATF1) upon TKO compared with NTC in HEK293T. (G) Co-immunoprecipitation analysis of the interaction between GFP-tagged EWSR1::ATF1 (GFP-EA1) and V5-tagged TRIM11 in HEK 293T cells. Cell lysates were subjected to immunoprecipitation with an anti-V5 antibody or control IgG, followed by immunoblotting for GFP, V5, and ACTB. Whole cell lysate is used as input.

To identify the components of the UPS that direct the degradation of EWSR1::ATF1, we performed a reporter-based CRISPR/Cas9 knockout screen. HEK293T Cas9 cells carrying the pArtichoke-EWSR1::ATF1 were transduced with the BISON CRISPR single guide (sg) RNA library, which targets more than 700 components of the ubiquitin-proteasome system, including E1 and E2 enzymes, E3 ligases, deubiquitinating enzymes and subunits of the proteasome^16,17^. Cells displaying the highest and lowest 5% of the EWSR1::ATF1-eGFP/mCherry ratio, corresponding to stabilized and destabilized fusion protein, respectively, were sorted and sequenced for hit identification (Figure 1C, Supplementary Figure S1A). The E3 ligases TRIM11, RNF216 and the deubiquitinating enzyme USP7 were recovered among genes whose knockout increased the stability of EWSR1::ATF1 (Figure 1D, Supplementary Table 1). As USP7 is broadly pleiotropic^19^, we focused our inquiry for fusion-specific effectors by focusing on TRIM11 and RNF216. Validation experiment using individual sgRNAs revealed that although knockout of both E3 ligases increased the eGFP/mCherry ratio, the fusion enrichment was stronger and more consistent across sgRNA for TRIM11 (Supplementary Figure S1B), which was therefore prioritized for further investigation.

To further interrogate the specificity of TRIM11 on EWSR1::ATF1, we assessed the effect of TRIM11 depletion on EWSR1::FLI1. The eGFP/mCherry ratios for EWSR1::FLI1 were unaffected by TRIM11 depletion, as were those of FLAG, whereas EWSR1::ATF1 levels strongly increased, indicating substrate specificity of TRIM11 towards EWSR1::ATF1 (Figure 1E). Immunoblotting confirmed the enrichment of EWSR1::ATF1 upon TRIM11 depletion without altering the levels of endogenous wild-type EWSR1 or ATF1, both in HEK293T cells (Figure 1F) and CCS cell line CCS210 (Supplementary Figure 1C), identifying EWSR1::ATF1 as a neo-substrate of TRIM11. To determine if TRIM11 associates with its substrate, we performed co-immunoprecipitation assays in HEK293T cells. Ectopically expressed GFP-tagged EWSR1::ATF1 interacted with V5-tagged-TRIM11 proteins, as demonstrated by the enrichment of V5-TRIM11 in the GFP-pull-down fraction and in the reciprocal setting EWSR1::ATF1-GFP in the V5-pull-down fraction (Figure 1G). To exclude any context specific factors, we also performed this assay in CCS210 cells (Supplementary Figure 1D), which validated the interaction. Together, these results established TRIM11 as a key regulator of EWSR1::ATF1 stability.

### High level of EWS::ATF1 supresses CCS proliferation

Next, we investigated the role of TRIM11 in stabilizing EWSR1::ATF1 in a CCS specific context. Single-guide RNA-mediated TRIM11 depletion was successfully achieved in all four models, with minor variation between sgRNAs and cell lines, which were all associated with increased EWSR1::ATF1 protein accumulation (Figure 2A). This increase was accompanied by impaired proliferation in all CCS cell lines, with stronger anti-proliferative effect in CCS210 and CCS-SN1 cells, compared with CCS24 and SUCCS1 cells. In contrast, TRIM11 knockout had no effect on proliferation of Ewing sarcoma cell line A673 or immortalized mesenchymal cell line SCP-1 (Figure 2B). These findings indicate that CCS cells exhibit a selective dependency on TRIM11-mediated regulation of EWSR1::ATF1 levels.

**Figure 2.**
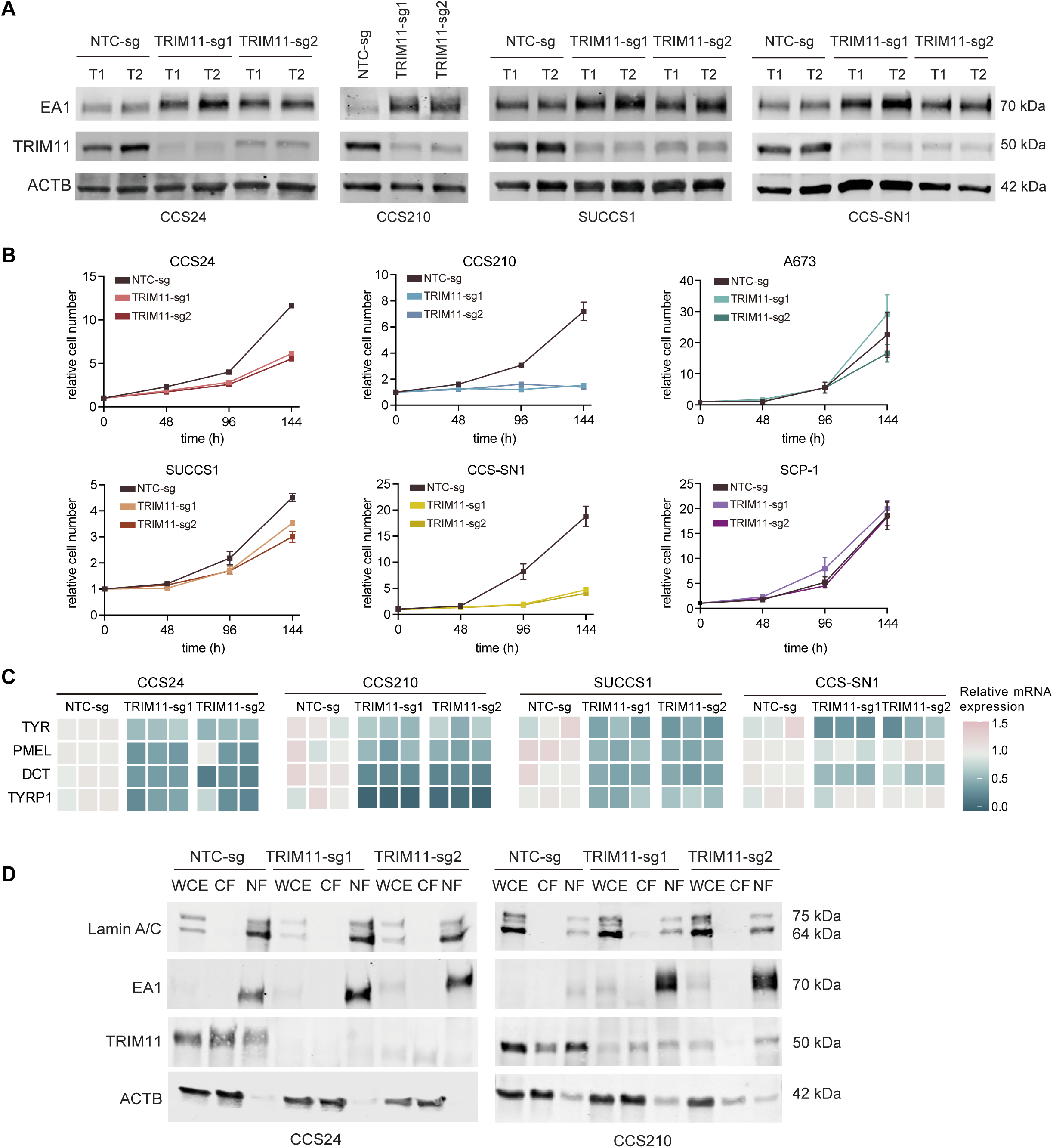
Higher level of EWSR1::ATF1 is unfavorable for CCS proliferation. (A) Western blot analysis of EWSR1::ATF1 (EA1) protein abundance following TRIM11 knockout with two independent single-guide RNAs (TRIM11-sg1 and TRIM11-sg2) in four CCS cell lines at two time points (T1 and T2). Samples were collected at day 4 (T1) and day 8 (T2) after puromycin selection. For CCS210, only T1 is shown because extensive cell death at T2 precluded the collection of sufficient material for analysis. (B) Growth curves of four CCS cell lines and two non-CCS cell lines (SCP-1 and A673) with TKO and non-targeting control (NTC). (C) qPCR analysis of MITF downstream target genes (TYR, PMEL, DCT, and TYRP1) in NTC and TKO cells. Results from three independent biological replicates are shown. (D) Cell fractionation followed by western blot analysis showing nuclear enrichment of EA1 upon TKO in CCS24 and CCS210. WCE, whole-cell extract; CF, cytoplasmic fraction; NF, nuclear fraction.

Previous studies have established that EWSR1::ATF1 hijacks the existing MITF transcriptional network to establish the oncogenic program of CCS while constraining neural crest and melanocytic differentiation^9^. Accumulation of a protein beyond the capacity of the cellular quality control machinery can produce aggregates that are toxic irrespective of the protein’s normal function^20^. To determine whether the loss of proliferation reflected this form of proteotoxicity or hyperactivation of the fusion protein, we first examined the MITF target genes, which is a key transcriptional program controlled by EWSR1::ATF1 after TRIM11 depletion. We observed that TRIM11-depletion mediated accumulation of EWSR1::ATF1 suppressed canonical MITF target genes involved in melanocytic differentiation, including TYR, PMEL, DCT and TYRP1 (Figure 2C). These data imply a transcriptional shift away from a differentiated melanocytic state and support a hyperactive state of EWSR1::ATF1 upon TRIM11 depletion. Cell fractionation showed that the accumulated EWSR1::ATF1 was present in the nuclear fraction (Figure 2D), confirming the localization of the surplus fusion protein at its site of transcriptional action, thereby adding to the evidence that the phenotype arises from excessive fusion-driven transcriptional activity. Collectively, these results implicate TRIM11 as a regulator of EWSR1::ATF1 stability, and its loss is leads to modulation of EWSR1::ATF1-driven transcriptional programs and impairs the viability of CCS cells.

### PCK1 is a downstream target of fusion hyperactivation

To mechanistically dissect how fusion hyperactivation restricts CCS growth, we profiled the proteome of TRIM11-depleted CCS24, CCS210 and SUCCS1 (Figure 3A, Supplementary Table 2). In agreement with the qPCR data (Figure 2C), the melanocytic differentiation markers PMEL and DCT were reduced when compared to proteomes of the cells transduced with non-targeting control sgRNA (Figure 3A). CDK2 was reduced in all three lines, in line with the loss of proliferation after TRIM11 depletion. Eight proteins increased in all three lines; the metabolic enzymes PCK1, POR and CTH, the membrane and structural proteins MYOF and TJP1, the signaling regulators RRAS2 and PTGES, and the deubiquitinating enzyme USP11. In support of these findings, examination of publicly available data from Ozenberger *et al*^10^ revealed modulation of these genes at the transcript level upon depletion of EWSR1::ATF1 (Figure 3B). Expression of these genes changed concordantly with EWSR1::ATF1 abundance in both directions, supporting their use as candidate readouts of fusion activity.

**Figure 3:**
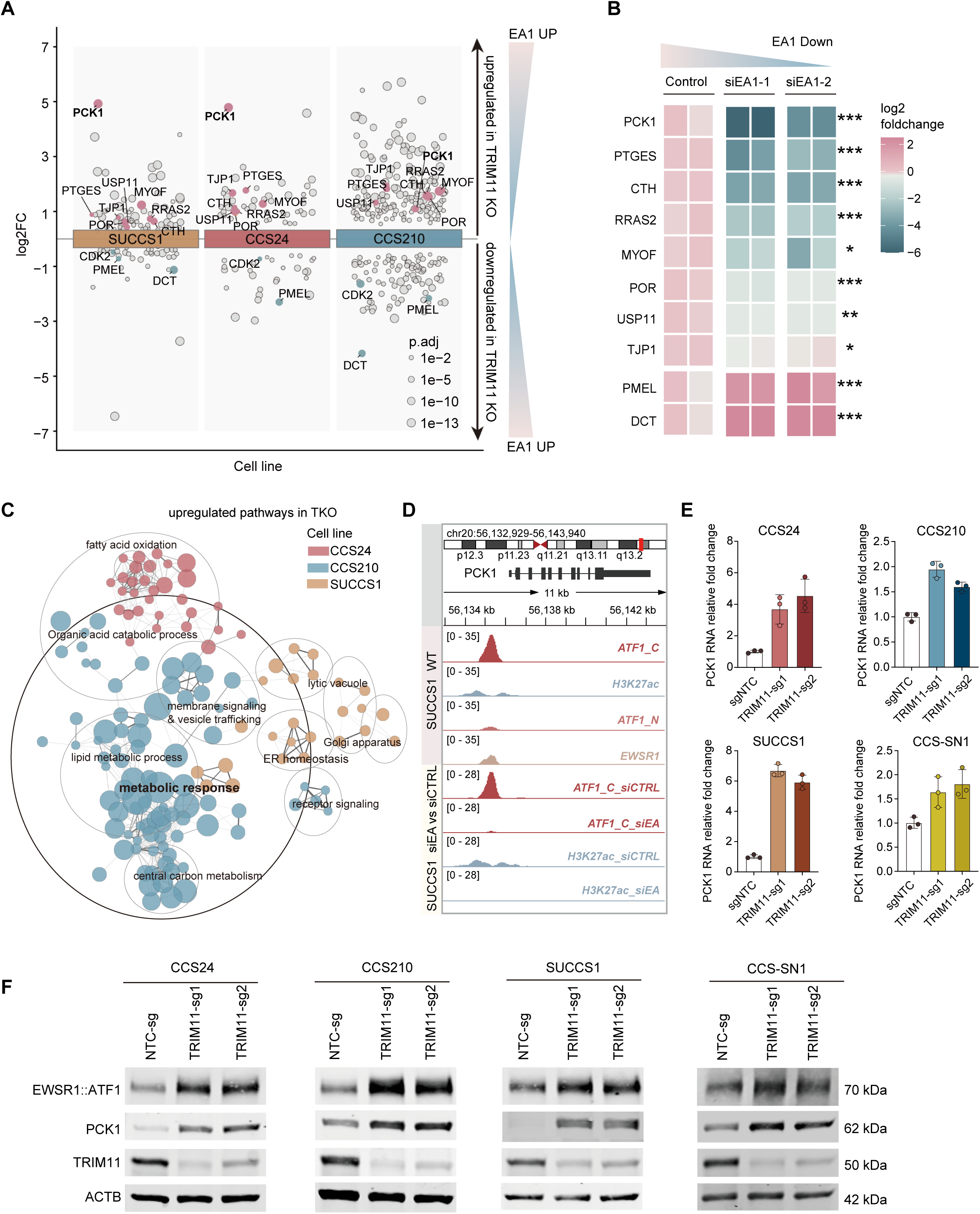
PCK1 is a downstream target of fusion hyperactivation. (A) Grouped strip plot showing upregulated and downregulated proteins in TRIM11 knockout (TKO) compared with non-targeting control (NTC). Only proteins that were significantly altered in both sgRNAs (p.adj < 0.05) are shown. Labeled proteins represent commonly upregulated or downregulated proteins identified across multiple cell lines. sgRNA, single-guide RNA, EA1: EWSR1::ATF1. (B) Heatmap showing relative expression changes of the indicated genes in SUCCS1 treated with siEA1 compared with control. Colors represent log2-transformed ratios calculated from normalized CPM values relative to the mean NTC expression. Stars indicate statistical significance from DESeq2 differential expression analysis (*adjusted P < 0.05, **adjusted P < 0.01, ***adjusted P < 0.001). CPM, counts per million. siEA1-2,siEA1-2: siRNA mediated EWSR1::ATF1 depletion. (C) Network analysis of upregulated pathways of TKO in three CCS cell lines. Each node represents one enriched Gene Ontology (GO) pathway (FDR < 0.05, minimum gene count >= 4) and is colored by cell line. Edge weight represents the Jaccard similarity between the protein sets contributing to the two GO terms/pathways. (D) ChIP-seq tracks showing EWSR1::ATF1 occupancy at the PCK1 proximal enhancer region. WT, wild type; siCTRL, control siRNA. siEA1: siRNA mediated EWSR1::ATF1 depletion. ATF1_C detects the C-terminal region shared by EWSR1::ATF1 and wild-type ATF1, ATF1_N detects the N-terminal region present only in wild-type ATF1. The EWSR1 track detects the N-terminal region of EWSR1 shared by wild-type EWSR1 and the EWSR1::ATF1 fusion protein. (E) qPCR analysis of PCK1 mRNA expression in four CCS cell lines following TRIM11 KO compared with NTC; n = 3 biological replicates. (F) Western blot of PCK1 protein expression in TKO and NTC samples across four CCS cell lines.

Pathway network analysis comparing control and TRIM11-depleted cells revealed metabolic processes as the dominant feature of the gene sets in all three cell lines (Figure 3C). The identity of the contributing proteins varied partly between the lines which resulted in cell-line specific but interconnected nodes, suggesting intrinsic heterogeneity between CCS cell lines. Despite this, metabolic processes were consistently enriched, indicating a shared response to fusion hyperactivation. Against this heterogeneity, PCK1 was the most strongly enriched protein in all three lines and was present in the majority of independent metabolic subnetworks (Supplementary Table 3). These results placed metabolic rewiring downstream of the increase in EWSR1::ATF1 abundance and identify PCK1 as its most consistent component.

To assess the potential role of EWSR1::ATF1 in regulating *PCK1*, we examined chromatin immunoprecipitation-sequencing data from the Möller *et al* study^9^ performed in CCS cells, in which EWSR1::ATF1 binding sites were mapped in the presence and absence of the fusion protein. At the *PCK1* locus, EWSR1::ATF1 occupied a site around 1 kb upstream of the transcription start site, concurrent with an H3K27ac signal. Both signals were markedly reduced following EWSR1::ATF1 depletion, consistent with direct regulation of PCK1 by the fusion protein. (Figure 3D). These data suggest that PCK1 is a direct transcriptional target of EWSR1::ATF1, consistent with its strong modulation upon fusion hyperactivation. These findings were further supported by upregulation of *PCK1* mRNA upon TRIM11 knockout, indicating transcriptional activation (Figure 3E), which was accompanied by PCK1 protein enrichment (Figure 3F), validating the findings from proteomics experiments in four CCS cell lines. Collectively, these findings identify PCK1 as a robust EWSR1::ATF1-responsive gene that is consistently induced following TRIM11 depletion.

### TRIM11 depletion induces TCA cataplerosis

Next, we sought to mechanistically dissect the impact of PCK1 protein enrichment in CCS cells. The tricarboxylic acid (TCA) cycle, a central metabolic hub that provides both energy and biosynthetic precursors, is maintained through a balance between carbon influx, termed as anaplerosis, and carbon efflux, called cataplerosis^21^. *PCK1* encodes cytosolic phosphoenolpyruvate carboxykinase (also abbreviated PEPCK), which catalyzes the conversion of oxaloacetate (OAA) to phosphoenolpyruvate (PEP), thereby redirecting the carbon flux from the TCA cycle toward gluconeogenic and related biosynthetic pathways, such as glyceroneogenesis and serine biosynthesis^22^. Thus, an increase in PCK1 abundance has the potential to alter carbon distribution in the cell, provided that the excess enzyme is catalytically active. Therefore, we first assessed PCK1 enzymatic activity in control and TRIM11-depleted cells. PCK1 activity was increased following TRIM11 depletion in CCS24, CCS210, and SUCCS1 cells (Figure 4A). Next, to underpin the potential metabolomic consequences of PCK1 enrichment, quantified intracellular levels of TCA cycle intermediates by targeted mass spectrometry. OAA, the substrate of PCK1, was reduced in all three cell lines, accompanied by reductions in the TCA cycle intermediates succinate, fumarate, and malate (Figure 4B).

**Figure 4.**
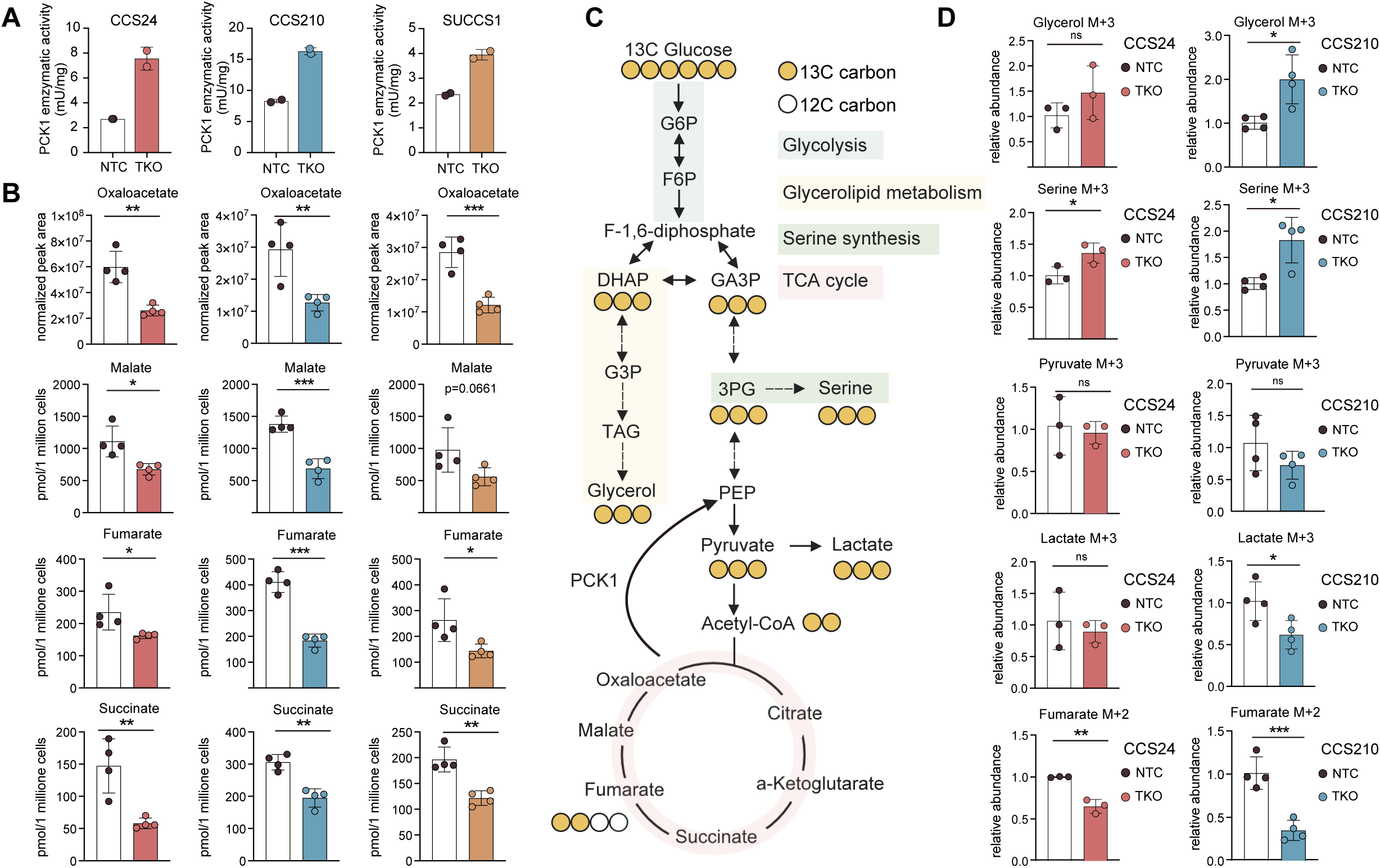
TRIM11 depletion induces TCA cataplerosis. (A) PCK1 enzymatic activity of CCS24, CCS210, and SUCCS1 in TRIM11 knockout (TKO) and non-targeting control (NTC). n= 2 technical replicates. Data is representative of two independent experiments. (B) Abundance of tricarboxylic acid (TCA) cycle intermediates in TKO and NTC cells of CCS24, CCS210, and SUCCS1. Calibrated metabolites are presented as concentrations. As no calibration curve was applied for oxaloacetate, it is presented as peak area normalized to the input cell number. Comparisons between two groups were performed using unpaired two-tailed Student’s t-tests (*P < 0.05, **P < 0.01, ***P < 0.001). (C) Schematic of 13C-glucose tracing; yellow-filled circles indicate incorporation of glucose-derived 13C in metabolites. G6P, glucose 6-phosphate; F6P, fructose 6-phosphate; DHAP, dihydroxyacetone phosphate; GA3P, glyceraldehyde 3-phosphate. G3P, glycerol 3-phosphate; 3PG, 3-phosphoglycerate; PEP, phosphoenolpyruvate; TAG, triacylglycerol. (D) Relative abundance of 13C-labeled fumarate, pyruvate, lactate, glycerol, and serine in TKO compared with NTC in CCS24 (n = 3) and CCS210 (n = 4). Comparisons between two groups were performed using unpaired two-tailed Student’s t-tests (*P < 0.05, **P < 0.01, ***P < 0.001).

To determine whether the depletion of TCA cycle intermediates was accompanied by carbon redistribution, we performed ^13^C-glucose isotope tracing in CCS24 and CCS210 cells following TRIM11 depletion. The analysis identified metabolite isotopologues (M+2, M+3) and their relative abundance. M+3 pyruvate levels remained unchanged in both lines in control and TRIM11-depleted conditions, while M+3 lactate was reduced in CCS210 and unchanged in CCS24, indicating preserved glycolytic carbon supply for entry into the TCA cycle (Figure 4C, D). M+2 fumarate was reduced in both lines, whereas M+3 serine was raised in both lines and M+3 glycerol in CCS210. These changes, together with lower TCA-cycle intermediates, support an altered central carbon metabolism upon TRIM11 depletion, with PCK1-mediated cataplerosis providing a mechanistic explanation.

### TRIM11 depletion reduces cellular respiration and damages mitochondria

Next, we sought to investigate whether these metabolic changes were accompanied by altered mitochondrial function. OXPHOS is tightly coupled to the TCA cycle, and intact OXPHOS activity relies on balanced TCA cycle homeostasis. Excessive cataplerotic flux can reduce the availability of TCA cycle intermediates, limiting substrate availability for mitochondrial respiration^23^.

To investigate whether cellular energy metabolism is altered with elevated cataplerosis activity in TRIM11 knockout cells, we used extracellular flux analysis to measure the oxygen consumption rate (OCR) and the extracellular acidification rate (ECAR). Depletion of TRIM11 lowered the OCR in all four CCS cell lines, with reductions in basal, maximal and ATP-linked respiration (Figure 5A to 5D). The extracellular acidification rate was unchanged in SUCCS1 and reduced in the other three lines (Figure S2A to S2D), and the energy map placed all four depleted lines closer to the quiescent state than their controls (Figure 5E). Estimated ATP production rates lowered (Figure S2E) and intracellular ATP abundance was reduced in all four lines (Figure S2F). We next examined whether impaired OXPHOS following TRIM11 loss was accompanied by mitochondrial damage, for which we assessed at the level of membrane potential, redox state and ultrastructure. JC-1 staining showed a lower aggregate to monomer ratio in TRIM11-depleted CCS24 and CCS210, indicating loss of mitochondrial membrane potential (Figure 5F). Mitochondrial superoxide measured by MitoSOX and total cellular reactive oxygen species measured by H2DCFDA were increased (Figure 5G, 5H), consistent with oxidative stress. Transmission electron microscopy further revealed altered mitochondrial ultrastructure, with TRIM11-depleted cells displaying mitochondria with widened intra-cristae spaces (black arrow) and enlarged crista-free regions of the matrix (asterisks, Figure 5I), providing morphological support for mitochondrial dysfunction^24^. Collectively, these results suggest that the respiratory defect following TRIM11 loss is accompanied by functional and structural mitochondrial damage (Figure 5J). To determine whether enhanced cataplerotic carbon withdrawal contributes to the altered energy metabolism, we supplemented cells with the anaplerotic substrate dimethyl-α-ketoglutarate (DM-α-KG). Indeed, the supplementation restored basal and maximal respiration, nearly completely and partially, in CCS24 and CCS210, respectively, towards control levels (Figure 5K), as well as reduced the fraction of annexin V positive cells in the depleted lines (Figure S2G). Finally, to assess if PCK1 could directly regulate the energy metabolism in CCS, we transiently overexpressed PCK1 in two CCS cell lines, CCS24 and CCS210 (Figure S2H), Ectopic PCK1 expression reduced oxidative phosphorylation (Figure S2J), phenocopying the direction of the respiratory defect induced by TRIM11 knockout, whereas PCK1 depletion (Figure S2I) enhanced mitochondrial respiration (Figure S2K). These complementary perturbations support a direct role for PCK1 in regulating mitochondrial bioenergetics in CCS. Taken together, these findings support a mechanistic model in which,by means of excessive PCK1 induction, TRIM11 loss increases cataplerosis, impairing mitochondrial oxidative metabolism and suppressing cell growth (Figure 6).

**Figure 5.**
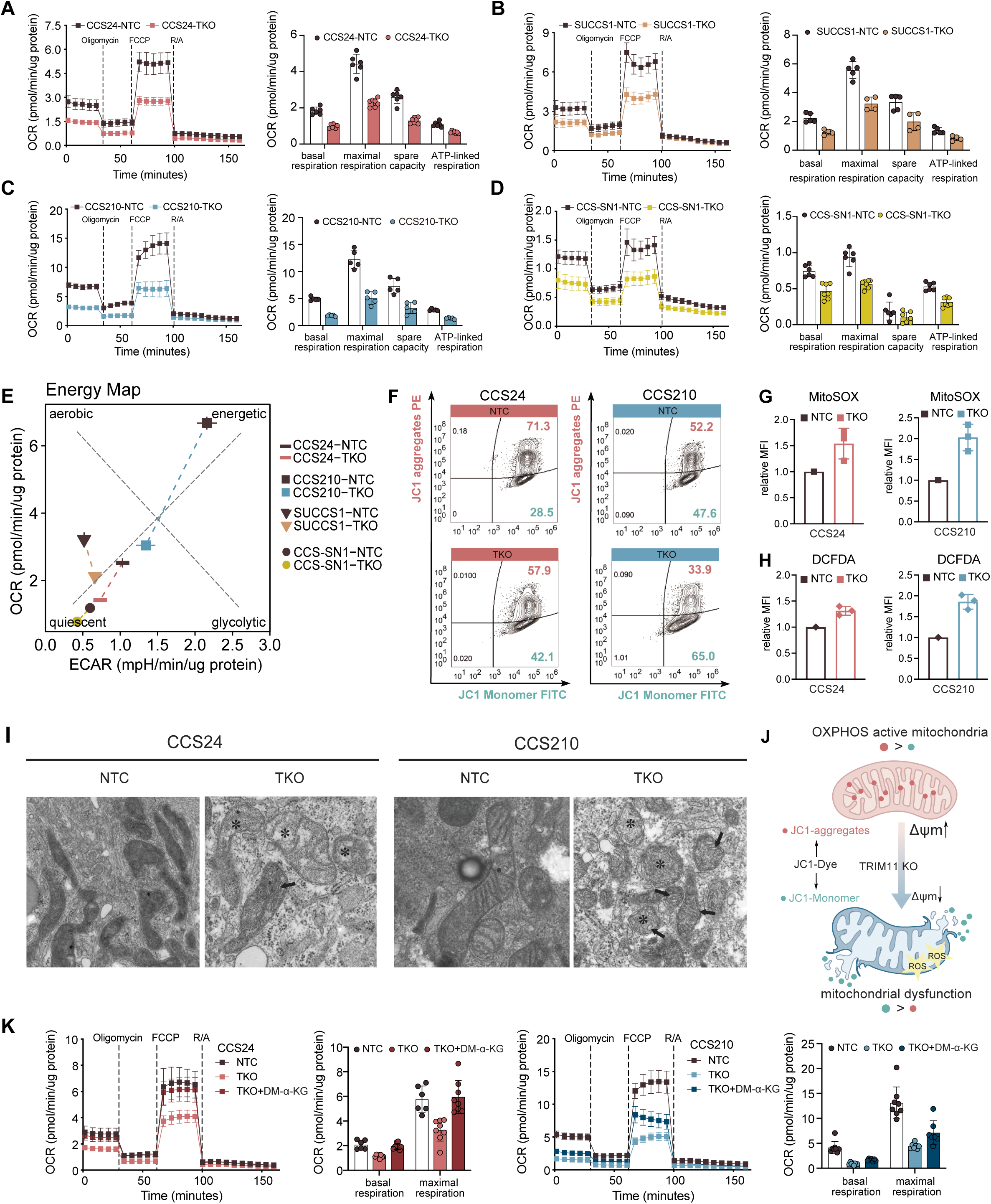
TRIM11 knockout reduced cellular respiration with mitochondrial damage. (A-D) Seahorse analysis of the oxygen consumption rate (OCR) in all four clear cell sarcoma (CCS) cell lines with non-targeting control (NTC) and TRIM11 knockout (TKO). Vertical dashed lines indicate the sequential addition of the indicated compounds. FCCP, carbonyl cyanide-p-trifluoromethoxyphenylhydrazone; R/A, rotenone/antimycin A. (E) Energy phenotype map summarizing alterations in cellular energy metabolism following TKO. ECAR, extracellular acidification rate. (F) Flow cytometric analysis of mitochondrial membrane potential detected by JC-1; data are representative of three biological replicates. (G) Flow cytometric analysis of mitochondrial reactive oxygen species (ROS) detected by MitoSOX. MFI, mean fluorescence intensity; n = 3 biological replicates. (H) Flow cytometric analysis of intracellular ROS detected by DCFDA; n = 3 biological replicates. DCFDA, 2′,7′-dichlorofluorescin diacetate. (I) Transmission electron microscopy images show mitochondrial ultrastructure in NTC and TKO cells of CCS24 and CCS210. Black arrors indicate distended cristae and asterisks indicate enlarged crista-free regions within the mitochondrial matrix. (J) Schematic illustration summarizing the mitochondrial alterations in TKO. (K) Seahorse analysis of the OCR in CCS24 and CCS210 with NTC, TKO, and TKO rescue with dimethyl α-ketoglutarate (DM-α-KG). All Seahorse results are representative of at least two independent experiments.

**Figure 6.**
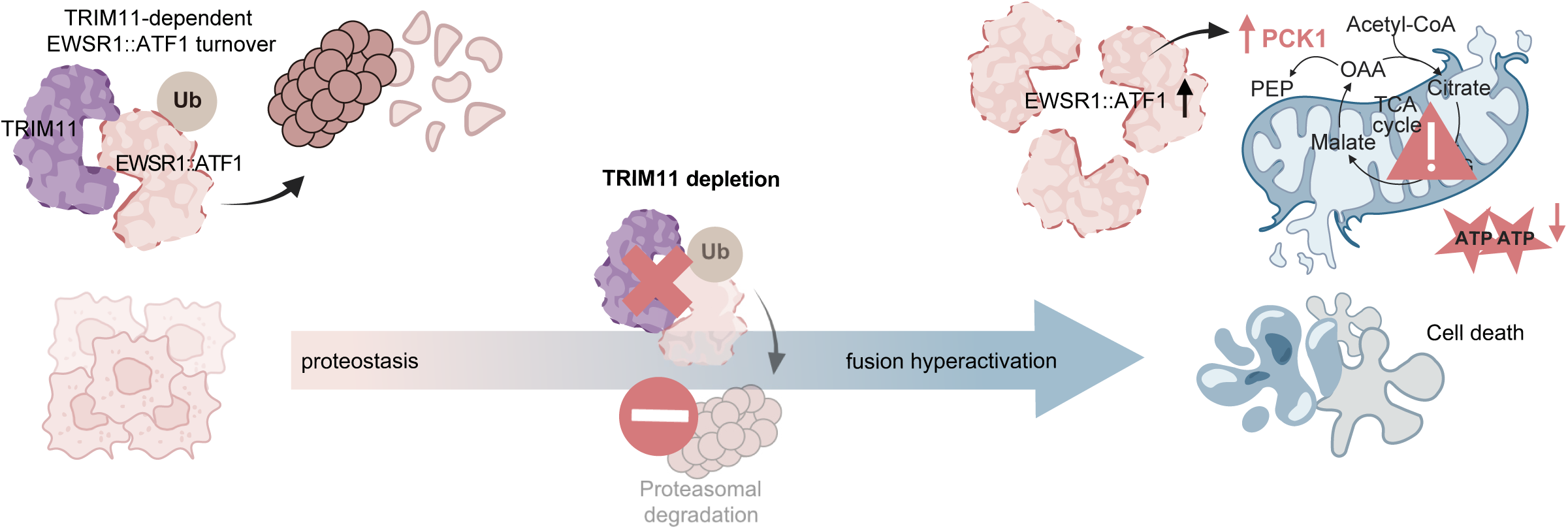
Proposed model of TRIM11-mediated regulation of EWSR1::ATF1 stability and metabolic homeostasis in CCS. TRIM11 depletion leads to EWSR1::ATF1 accumulation and hyperactivation, resulting in excessive PCK1 induction and contributing to increased TCA-cycle cataplerosis and depletion of the TCA-cycle intermediate pool. The resulting metabolic state is characterized by impaired oxidative phosphorylation, ATP depletion and mitochondrial dysfunction, together with reduced CCS cell proliferation and viability. OAA, oxaloacetate; PEP, phosphoenolpyruvate; TCA, tricarboxylic acid.

### PCK1 expression is associated with aggressive clinical features in CCS

Having identified PCK1 as a fusion target and linked its induction to dysregulated mitochondrial metabolism following TRIM11 depletion, we next asked whether its level of expression is itself informative of disease behavior in patients. To this end, we analyzed RNA-seq data of CCS tumors (n=29) and correlated PCK1 expression levels with patient outcome (Figure 7A, Supplementary Table 4). High PCK1 expression was associated with shorter progression free survival (13.77 months) compared to patients with low PCK1 expression (55.98 months) in patients diagnosed with primary disease (n=19). In the full cohort, high PCK1 expression correlated with shorter overall survival (15.36 months) than patients with low PCK1 expression (60.81 months). Moreover, patients presenting with metastatic disease at diagnosis exhibited higher PCK1 expression than those who developed disease progression after initial diagnosis (Figure 7B). Transcriptomic correlation analysis revealed PCK1 expression is positively correlated with gene programs related to cell proliferation and invasion (Figure 7C, Supplementary Table 5). Among individual genes, PCK1 expression positively correlated with SNAI1, which is a key regulator of invasive and migratory programs as previously introduced, and negatively correlated with MTSS1, a metastasis suppressor that is frequently lost in advanced tumors (Figure 7D). Together, these findings suggest that PCK1 may serve as a prognostic marker for CCS.

**Figure 7.**
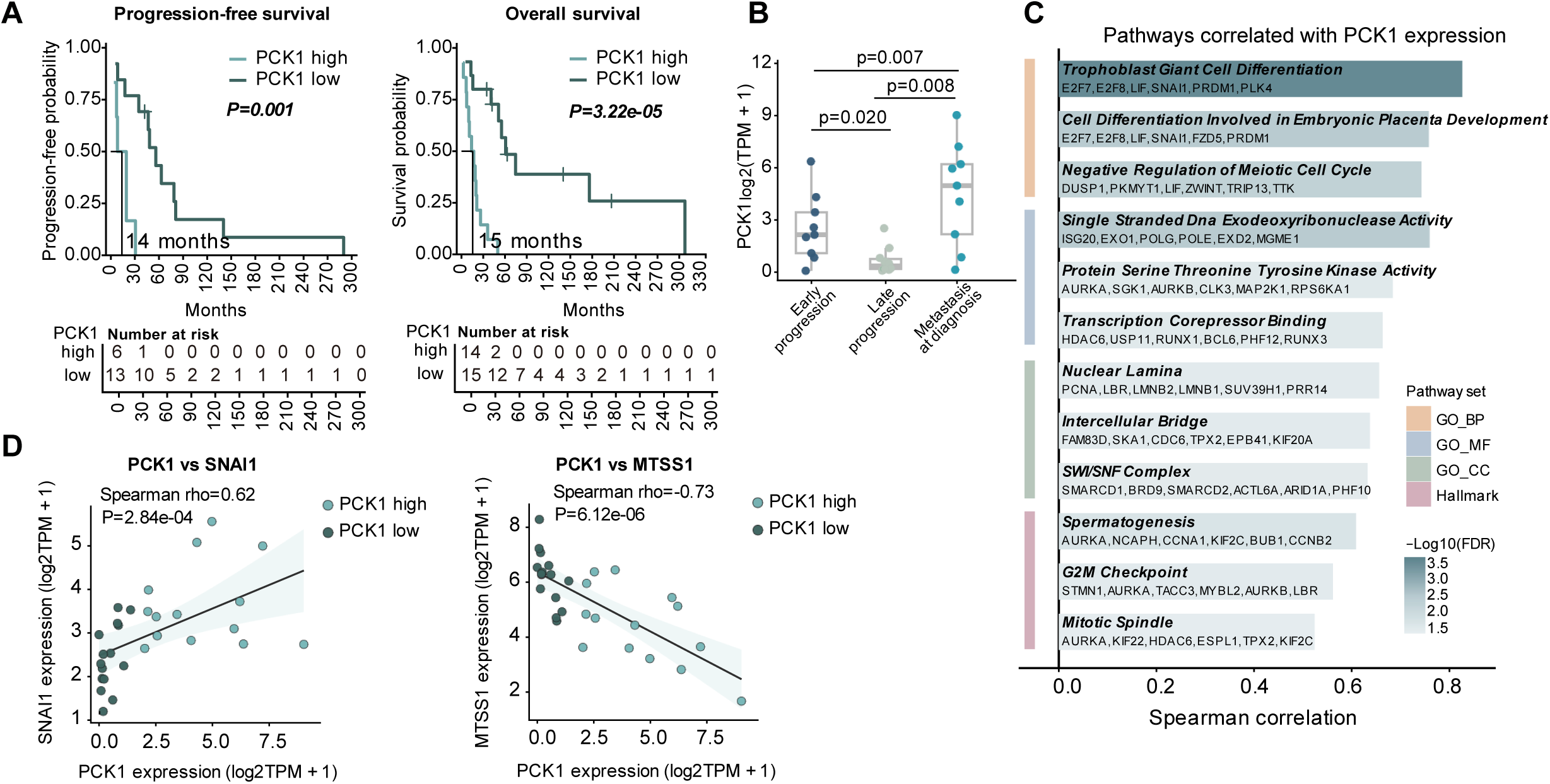
PCK1 is associated with aggressive clinical features in CCS. (A) Kaplan-Meier curves show progression-free survival (PFS; n = 19 primary cases without metastasis at diagnosis) and overall survival (OS; n = 29) stratified by median PCK1 expression. P values were calculated using the log-rank test. Censored cases are indicated by tick marks. The indicated 14- and 15-month values denote the median PFS and OS, respectively, of the PCK1-high group. The number-at-risk tables report the numbers of patients in each PCK1-expression group who remained under observation and event-free at each corresponding time point. (B) Boxplot showing PCK1 expression across progression groups defined in the clinical annotation. Early and late progression were classified using the median PFS time of primary cases without metastasis at diagnosis, ∼ 900 days. The metastasis-at-diagnosis group included patients with metastatic disease present at initial diagnosis. Right-censored cases were excluded from this comparison. Overall group differences were assessed by Kruskal–Wallis test, with pairwise Wilcoxon rank-sum tests shown above the indicated comparisons. (C) Pathway barplot shows the top PCK1-correlated GO and Hallmark pathways; bar length represents Spearman correlation with PCK1, and color indicates −log10(FDR). (D) gene expression correlation between PCK1 and SNAI1 or MTSS1 in CCS tumor RNA-seq data.

## Discussion

Our study identified TRIM11, an E3 ligase, as a regulator of the stability of EWSR1::ATF1 in CCS. We show that depletion of TRIM11 increases fusion protein abundance, and in turn, amplifies fusion-driven transcriptional programs, accompanied by metabolic rewiring in which PCK1 emerged as a responsive component. This metabolic state was characterized by increased cataplerosis with depletion of the TCA cycle intermediate pool with impaired oxidative phosphorylation and reduced cellular proliferation. These findings position TRIM11 depletion as a therapeutic approach in CCS and identify PCK1 as the effector contributing to this altered central carbon metabolism.

Oncogenic signaling is sustained not only by the mechanisms that activate driver pathways, but also by the regulatory processes that constrain their activity below a toxic threshold. In RTK-RAS-driven lung cancer, disruption of negative feedback regulators produced lethal ERK hyperactivation, showing that factors attenuating oncogenic signaling may themselves constitute cancer dependencies^25^. A comparable dosage constraint has been described for EWSR1::FLI1 in Ewing sarcoma, where proteasomal turnover mediated by TRIM8 prevents accumulation of the fusion protein beyond the tolerated level^18^. Our findings extend this principle to EWSR1::ATF1 in CCS and suggest dosage control as a shared requirement of the two EWSR1-rearranged sarcomas. Both fusion proteins are turned over through the ubiquitin-proteasome system by a RING E3 ligase of the TRIM family, yet each depends on a distinct member, since TRIM11 depletion did not lead to enrichment of EWSR1::FLI1 (Figure 1E) and the proliferation capacity of cell line driven by EWSR1::FLI1 also remained unaffected (Figure 2B). The consequences of ligase depletion are, in turn, determined by lineage and the oncogenic transcriptome that each fusion protein orchestrates. Systematic assessment of further EWSR1-rearranged fusions may therefore establish dosage control as a common denominator of this group of cancers, while mechanistic dissection of their individual downstream programs may uncover further biology and genotype-selective vulnerabilities.

The identification of TRIM11 as an E3 ligase governing EWSR1::ATF1 turnover, together with the lethality of depletion in CCS cell lines, suggests two preclinical therapeutic avenues. The first is to enhance degradation of the driver fusion protein by inducing proximity between TRIM11 and EWSR1::ATF1. Because our data identify EWSR1::ATF1 as a neosubstrate of TRIM11, a “molecular glue” that stabilizes this interaction could selectively promote fusion protein degradation in translocation-positive cells, potentially offering greater specificity than conventional transcription-factor inhibition. Development of such compounds has historically proven demanding, and the intrinsically disordered character of the EWSR1 portion presents an additional obstacle to defining a stable interface for structure-guided design. The second avenue would be exploiting the lethality of TRIM11 depletion by pharmacological inhibition. Unaltered cell viability in fusion-negative cell lines support this therapeutic window. Crystal structures of the PRYSPRY domains of nine TRIM proteins, including TRIM11 (PDB: 7QS1) have been characterized. Of note, TRIM11 structure features a subdomain swap absent from the other family members together with variation in the loops flanking the canonical substrate-binding site^26^. This difference may provide the structural basis for TRIM11-selective binder design. Taken together, these considerations suggest that the TRIM11 and EWSR1::ATF1 axis is tractable from two distinct directions, namely targeted degradation of the fusion oncoprotein by induced proximity with TRIM11 by “molecular glue” degrader, or fusion hyperactivation by TRIM11 inhibition.

CCS has thus far been regarded as a uniform entity on the basis of single recurrent driver fusion oncoprotein, however, our data reveal heterogeneity in cellular response to TRIM11 depletion. Proliferation was reduced in all four cell lines following TRIM11 depletion, although the magnitude differed, with CCS210 and CCS-SN1 more strongly affected than CCS24 and SUCCS1. Similarly, the proteomic response to TRIM11 depletion showed a comparable pattern of enriched metabolic processes in all cell lines (CCS24, CCS210 and CCS-SN1) investigated, while the contributing proteins differed such that each line formed its own subnetwork. Basal energy metabolism likewise differed between the lines prior to perturbation, with the energy map placing them at distinct positions along the respiratory and glycolytic axes. These findings suggest potentially clinically relevant heterogeneity between CCS lines that warrant further investigation. Despite this, PCK1 emerged across CCS models as a consistent fusion-responsive metabolic node, providing a potential link between fusion accumulation, altered carbon allocation and mitochondrial dysfunction.

An important aspect of cancer metabolism is the dynamic allocation of carbon between bioenergetic and biosynthetic pathways to accommodate fluctuating metabolic demands^27^. Rather than maximizing ATP production or biomass synthesis alone, proliferating tumor cells continuously balance carbon utilization between oxidative metabolism and anabolic processes to sustain both energy homeostasis and cell growth^28^. Positioned at this metabolic branch point, PCK1 governs the distribution of TCA cycle-derived carbon between mitochondrial oxidation and anabolic metabolism^29^. Consistent with this role, independent PCK1 perturbation experiments showed that increasing and decreasing PCK1 abundance altered mitochondrial respiration in opposite directions, supporting a direct role for PCK1 in regulating mitochondrial bioenergetics in CCS.

Adding to this experimentally defined metabolic function, patient outcome data point to the clinical relevance of PCK1. At endogenous levels, high PCK1 expression occurred in tumors with poorer clinical outcomes and with transcriptional programs associated with proliferation and invasion, supporting the potential prognostic relevance of PCK1. The association of endogenous PCK1 expression with aggressive disease appears paradoxical with the metabolic impairment accompanying its induction following TRIM11 loss. A possible explanation is that tumors with high PCK1 expression have adapted their metabolic program to accommodate sustained cataplerotic flux, such that PCK1 activity may not only remain compatible with the metabolic requirements of tumor proliferation, but even support metastatic processes, potentially through carbon redistribution into biosynthetic pathways. Its excessive induction to supraphysiological levels following TRIM11 loss may disrupt the balance, becoming detrimental when cataplerotic carbon withdrawal exceeds the capacity of anaplerotic pathways to sustain the TCA cycle. These findings position the axis comprising TRIM11, EWSR1::ATF1 and PCK1 as an actionable route in CCS pathobiology, with PCK1 serving both as the effector through which TRIM11 depletion imposes its phenotype, and as a candidate prognostic biomarker.

We acknowledge that our study has several limitations. First, our study does not investigate the TRIM11 substrate spectrum beyond EWSR1::ATF1. Identification of these substrates will help distinguish the consequences of EWSR1::ATF1 accumulation from other potential effects of TRIM11 depletion, which may nominate candidate secondary dependencies for combination with TRIM11 inhibition. Second, our findings are based on genetic depletion in cell line models, since pharmacological TRIM11 inhibitors are not yet available. Their development, together with suitable *in vivo* models, would allow oncogene hyperactivation to be examined *in situ*, allowing study of drug penetration and oncogene doasge. Third, our cohort of 29 patients, although also sizeable for an ultra-rare sarcoma, yet remains too small to determine whether PCK1 carries prognostic information independent of the clinical factors established in CCS^30^. Future studies in extended cohorts, together with development of a clinically applicable assay, will help establish the utility of PCK1 in risk assessment. Furthermore, it remains to be clarified whether the correlation between high PCK1 expression and poor clinical outcomes is merely reflecting overall level of fusion-driven transcriptional activity, or whether it is directly related to modulation of bioenergetic and biosynthetic pathways downstream of PCK1. Follow-up studies evaluating the functional impact of PCK1-dependent metabolic pathways, including their effects on biosynthesis and tumor growth in CCS will help clarify the role of PCK1 in tumor progression in presence and absence of TRIM11 depletion.

In summary, this study demonstrates that TRIM11 regulates the abundance of EWSR1::ATF1 in CCS, and that release of this constraint is lethal through increased withdrawal of carbon from the TCA cycle and reduced cell viability that is mediated by excessive PCK1 induction. Our study establishes the TRIM11 - EWSR1::ATF1 - PCK1 axis as an entry point for therapeutic intervention in a disease that lacks effective conventional or molecularly directed treatment.

## Materials and methods

### Cell lines and cultivation

The clear cell sarcoma (CCS) cell lines CCS24, CCS210, were kindly provided by Jonathan Fletcher (Harvard Medical School and Mass General Brigham Hospital in Boston). SUCCS-1 (CRL-2971™) were purchased from the American Type Culture collection (ATCC). CCS-SN1 (HU-25.6) cell line was generated in the laboratory of Sebastian Bauer (University Hospital Essen, Essen, Germany) under the Human Sarcoma Model Initiative (https://www.husarc.org/). All CCS cell lines were cultured in RPMI 1640 supplemented with 15% fetal bovine serum (FBS), 1% L-glutamine, and 1% penicillin–streptomycin. Cas9-expressing HEK293T cells were provided by Mikołaj Słabicki (Broad Institute, Cambridge, MA, USA) and cultured in DMEM supplemented with 10% FBS, 1% L-glutamine, and 1% penicillin–streptomycin. SCP-1 cells were kindly provided by Thomas Kindler (University Cancer Center Mainz, Mainz, Germany) and cultured in MEM supplemented with 10% FBS, 1% L-glutamine, and 1% penicillin– streptomycin. A673 cells were kindly provided by Thomas Grünewald (German Cancer Research Center, Heidelberg, Germany) and cultured in RPMI 1640 supplemented with 10% FBS, 1% L-glutamine, and 1% penicillin–streptomycin. All cell lines were maintained at 37°C in a humidified atmosphere containing 5% CO_2_, and were regularly tested for mycoplasma contamination. Cells were passaged by trypsinization when they reached 70-80% confluency.

### Plasmids, lentivirus production, and transduction

The EWSR1–ATF1 fusion construct was cloned into the pArtichoke lentiviral reporter vector (Addgene plasmid #73320), which encodes mCherry as an internal control and expresses protein of interest tagged with GFP. Lentiviral particles were produced in HEK293T packaging cells by co-transfecting the appropriate transfer vector with psPAX2 (Addgene plasmid #12260) and pMD2.G (Addgene plasmid #12259). HEK293T packaging cells were transfected using TransIT-LT1 reagent (Mirus Bio, #MIR2304). Virus-containing supernatants were harvested, filtered through 0.40-µm filters, aliquoted, and stored at −80°C for further use. Before screening, Cas9-expressing cells were maintained under blasticidin selection. After transduction, reporter cells were either selected with puromycin or enriched by flow cytometry, as appropriate.

### Cloning of overexpression constructs

The open reading frame (ORF) of PCK1 (ORFeome Collaboration OCAAo5051H08042) in gateway entry clone format or a multiple cloning site (MCS) as empty vector control was shuttled in the lentiviral expression vector rwpLX305-FLAG_GW-IRES-Puro (Cellular Tools Core Facility, DKFZ Heidelberg) by site specific recombination. The resulting expression construct drives the PKC1 expression from a CMV promoter and a Flag-Tag to the n-terminus of PKC1. For selection stable expressing cell lines the mRNA expression is coupled to an IRES-Puromycin selection marker.

### Cloning of single guide RNAs

For individual knockout experiments, complementary oligonucleotides encoding sgRNAs targeting *TRIM11* or *PCK1*, or a non-targeting control (*NTC*), were phosphorylated, annealed, and ligated into the BsmBI-digested pLKO5.sgRNA.EFS.PAC lentiviral vector (Addgene plasmid #57825). Candidate sgRNA sequences/protospacers were selected using CRISPick, the Broad Institute Genetic Perturbation Platform (GPP) web tool. The cloning oligonucleotide sequences are listed in the table below.

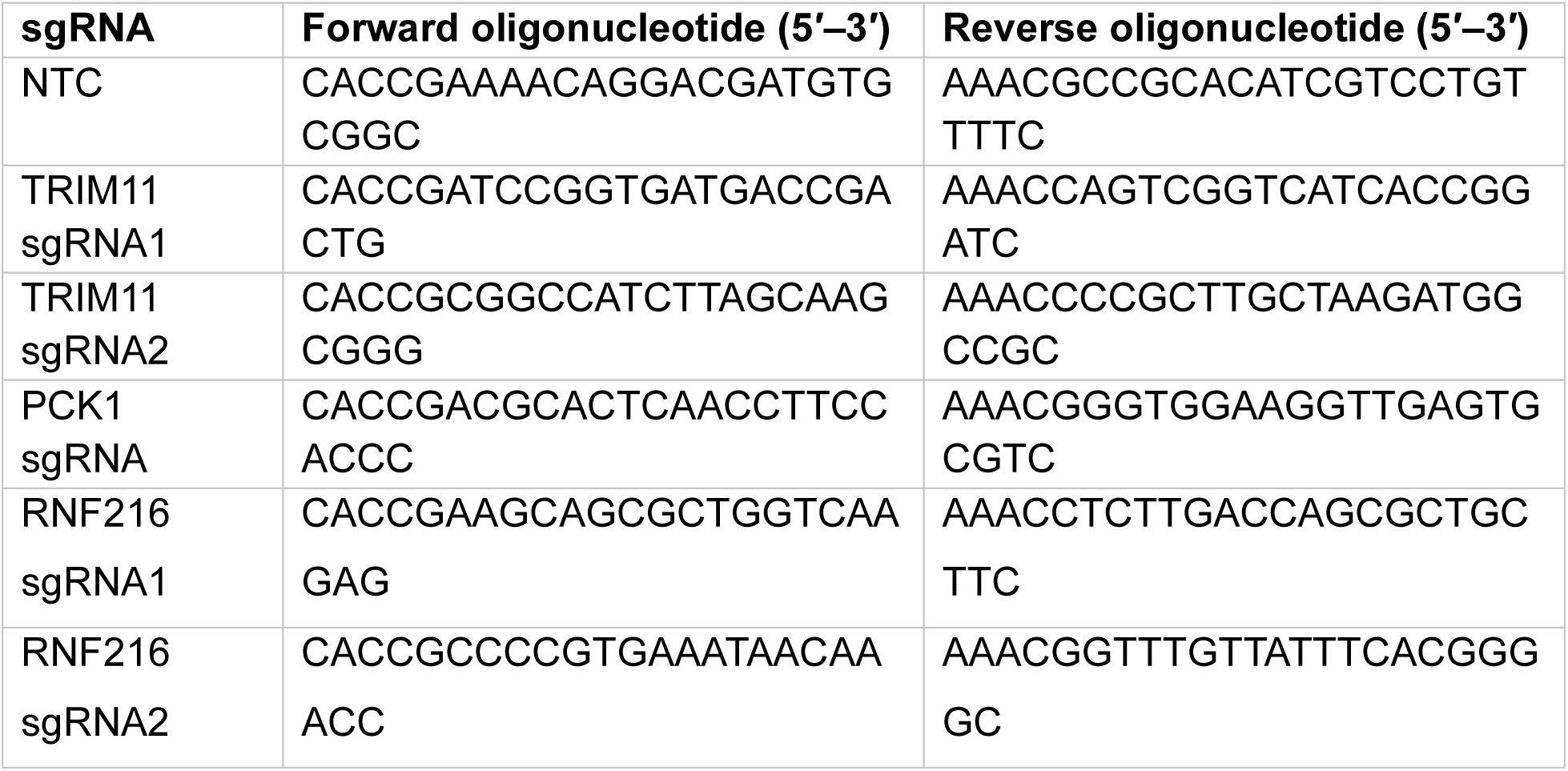

Ligation products were transformed into Stbl3 competent cells, plasmid DNA recovered from bacterial colonies was purified and sequenced were verified by Sanger sequencing using the U6 forward primer. The resulting sgRNA constructs were delivered to target cells by lentiviral transduction.

### Flow cytometric analysis of the EWSR1::ATF1 reporter

GFP and mCherry fluorescence signals were acquired on a FACSCelesta SORP cell analyzer (BD Biosciences) using FACSDiva software. Debris were excluded using forward- and side-scatter parameters, followed by exclusion of doublets using forward-scatter area and height. Events positive for both GFP and mCherry were selected, and the GFP:mCherry fluorescence ratio was calculated for each event in FlowJo (v10.2). For inhibitor experiments, cells were treated for 2 h with 0.5 µM MG132 or 1 µM MLN7243 before acquisition; DMSO-treated cells served as controls.

### Bison CRISPR–Cas9 knockout screen

The Bison sgRNA library was kindly provided by Mikołaj Słabicki. The library contains 2,852 sgRNAs targeting 713 components of the ubiquitin–proteasome system and control genes, with four sgRNAs per gene^16^. HEK293T Cas9 cells stably expressing the EWSR1::ATF1 pArtichoke reporter were transduced with pooled Bison lentivirus, with library representation maintained throughout the screen. Nine days after transduction, debris, dead cells, doublets, and cells lacking either reporter fluorophore were excluded. Among GFP- and mCherry-double-positive events, the upper and lower 5% of the GFP:mCherry fluorescence-ratio distribution were sorted using a FACSAria III cell sorter equipped with an 85-µm nozzle (BD Biosciences). Genomic DNA was isolated from sorted cells, sgRNA cassettes were amplified by two-step PCR, and the resulting libraries were sequenced on an Illumina NextSeq 550 system. Demultiplexed reads were trimmed, and sgRNA count tables were generated using MAGeCK version 0.5.9^31^. Within each sample, sgRNA counts were normalized to that sample’s mean count and then scaled to the global mean across samples. For each of three replicates, high-to-low count ratios were calculated and sgRNAs were ranked accordingly. sgRNA ranks were summed across replicates, and each gene was assigned the median of the resulting rank for its four sgRNAs. Empirical two-sided P values were estimated from a null distribution generated by 1,000 random permutations of sgRNA ranks and were adjusted for multiple testing using the Benjamini–Hochberg method. Genes with a Benjamini–Hochberg-adjusted P value (false discovery rate) below 0.05 were classified as screen hits.

### Western Blot

Cells were lysed in buffer containing 1% SDS, 10 mM Tris–HCl, 1 mM sodium orthovanadate, protease inhibitor cocktail (Thermo Fisher Scientific, #11834101), and phosphatase inhibitor cocktail (Thermo Fisher Scientific, #78420), and incubated with shaking in a thermoblock at 95°C for up to 20 min. Protein concentrations were determined using the Pierce BCA Protein Assay Kit (Thermo Fisher Scientific, #23225). Samples were mixed with NuPAGE LDS Sample Buffer (Invitrogen, #NP0007) and reducing agent (Invitrogen, #NP0009), heated at 95°C for 5 min, and separated on 10% Mini-PROTEAN TGX gels (Bio-Rad). Proteins were transferred to PVDF membranes and blocked in 5% non-fat milk in TBS containing 0.1% Tween-20 for 1 h at room temperature. Membranes were incubated overnight at 4°C with primary antibodies: anti-TRIM11 (Proteintech, #84651-3-RR; 1:2,000), anti-EWSR1 (Santa Cruz Biotechnology, #sc-28327; 1:500), anti-ATF1 (Santa Cruz Biotechnology, #sc-241; 1:500), anti-GAPDH (Cell Signaling Technology, #5174; 1:2,000), anti-lamin A/C (BioLegend, #600002; 1:500), anti-α-tubulin (Abcam, #ab7291; 1:3,000), anti-PCK1 (Cell Signaling Technology, #12940S; 1:2,000); 1:1000) anti-V5 (Cell Signaling Technology, #14793S; 1:1000), anti-GFP (Cell Signaling Technology, #2956S; 1:1000) anti-beta-Actin (Cell Signaling Technology, #4967S; 1:1000) and (Santa Cruz, #sc-47778; 1:1000). Membranes were washed three times with TBS-T and incubated for 1 h at room temperature with IRDye-compatible anti-mouse (Cell Signaling Technology, #5257S, #5470S; 1:10,000) or anti-rabbit secondary antibodies (Cell Signaling Technology, #5151S, #5366S; 1:10,000) emitting at 680 or 800 nm where ever applicable. Signals were acquired using an Odyssey CLx imaging system (LI-COR) and analyzed using Image Studio Lite and Fiji.

### Co-immunoprecipitation

Cells were transiently co-transfected with indicated expression constructs using TransIT-LT1 transfection reagent. At 48 h after transfection, cells were treated with 5 µM MG132 for 4 h before harvesting to inhibit proteasomal degradation. Cells were lysed in RIPA IP lysis buffer containing 50 mM Tris–HCl (pH 7.6), 150 mM NaCl, 1 mM EDTA, and 1% Triton X-100, supplemented with protease and phosphatase inhibitor cocktails. Equal amounts of total protein (3 mg) were incubated overnight at 4°C with the appropriate primary antibody (anti-GFP or anti-V5), followed by incubation with Protein A/G agarose beads (Santa Cruz, #sc-2003) for 3 h at 4°C. Beads were washed three times with wash buffer containing 50 mM Tris–HCl (pH 7.5), 150 mM NaCl, 2 mM EDTA, 1% Triton X-100, 1% sodium deoxycholate, and 0.1% SDS.

Bound proteins were eluted by boiling the beads for 5 min at 95°C in 2× SDS sample buffer consisting of NuPAGE LDS Sample Buffer (Invitrogen, #NP0007) and NuPAGE Sample Reducing Agent (Invitrogen, #NP0009). Samples were analyzed by SDS-PAGE and Western blotting using the Odyssey CLx Imaging System (LI-COR Biosciences). Whole-cell lysate (3% of the total lysate) was included as an input control. Mouse monoclonal, anti-rabbit IgG light chain specific (Abcam, #ab-99697) secondary antibody was used to prevent interference from heavy immunoglobulin chains in the binding assay.

### Cell fractionation

CCS24 (2 × 10⁶) or CCS210 (1 × 10⁶) cells were incubated for 15 min on ice in cytoplasmic lysis buffer containing 10 mM HEPES (pH 7.6), 0.1 mM EDTA, 10 mM KCl, 1 mM DTT, 50 mM NaF, 0.5 mM PMSF, 5% glycerol, and protease inhibitor cocktail. NP-40 was added to 1%, and samples were centrifuged at 1,000 × g for 5 min. The supernatant was collected as the cytoplasmic fraction. Nuclear pellets were washed three times with cytoplasmic lysis buffer and resuspended in nuclear lysis buffer containing 1% SDS, 10 mM Tris–HCl, 1 mM sodium orthovanadate, protease inhibitor cocktail, and phosphatase inhibitor cocktail. Samples were heated at 95°C for 10 min and centrifuged for 2 min at 13000 rpm and the supernatant was collected as the nuclear fraction. Protein concentrations were determined using BCA assay.

### Reverse transcription quantitative PCR

Total RNA was isolated using the RNeasy Plus Midi Kit (Qiagen, #75144) according to the manufacturer’s instructions. RNA concentration was measured using a NanoDrop spectrophotometer, and 2 µg RNA was reverse-transcribed using the High-Capacity cDNA Reverse Transcription Kit (Thermo Fisher Scientific, #10400745). Quantitative PCR (qPCR) was performed using iTaq Universal SYBR Green Supermix (Bio-Rad, #1725120) on a CFX96 Real-Time PCR Detection System (Bio-Rad). qPCR reactions were performed for 40 amplification cycles, followed by melting-curve analysis. Relative expression was calculated using the ΔΔCt method, with *PBGD* as the reference gene and normalized to the corresponding control condition.

Primer sequences:

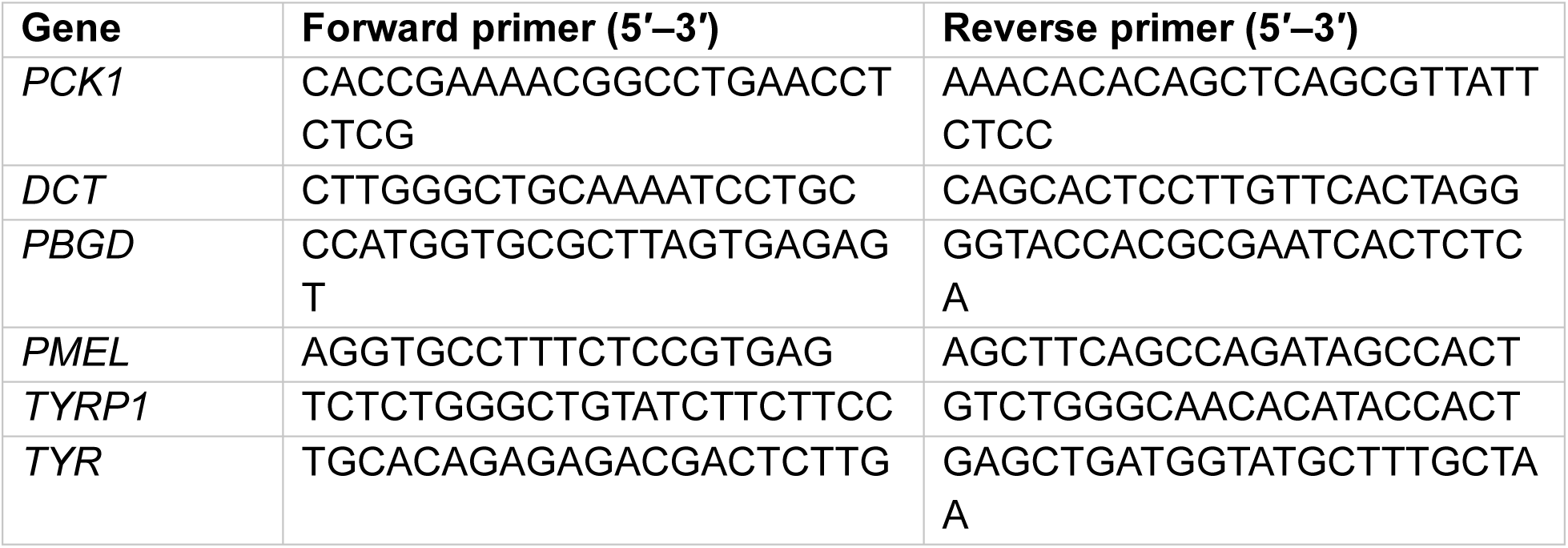

### Cell proliferation assays

Cells were seeded in 24-well plates and viable cell were counted from day 2 to day 6 after seeding. Cell suspensions were mixed with 0.4% trypan blue, and viable cells were quantified using a LUNA-II automated cell counter (Logos Biosystems).

### Full-proteome mass spectrometry

Total of 5ug proteins were digested with trypsin using an AssayMAP Bravo liquid-handling system (Agilent Technologies) running the automated single-pot solid-phase-enhanced sample preparation (autoSP3) protocol. A 5-µg peptide aliquot was reserved for full-proteome analysis. Dried peptides were reconstituted in 97.4% water, 2.5% hexafluoro-2-propanol, and 0.1% trifluoroacetic acid, and 500 ng peptide was analyzed per injection.

LC–MS/MS was performed using a Vanquish Neo UPLC system coupled directly to an Orbitrap Exploris 480 mass spectrometer (Thermo Fisher Scientific). Peptides were desalted on an Acclaim PepMap300 C18 trapping cartridge (5 µm, 300 Å) and separated on a nanoEase M/Z Peptide analytical column (75 µm × 200 mm, 1.7 µm, 300 Å; Waters) at 300 nl/min. Solvent A was 0.1% formic acid in water and solvent B was 0.1% formic acid in acetonitrile. Solvent B was increased from 4% to 30% over 135 min, followed by an increase to 80%, return to 2%, and re-equilibration. Data-dependent acquisition comprised a full scan at 120,000 resolution over m/z 380–1,400 (300% AGC target; 45-ms maximum injection time), followed by MS/MS acquisition for up to 2 s. Precursors were isolated with a 1.4-m/z window and fragmented at 26% normalized collision energy. Fragment spectra were acquired at 15,000 resolution (100% AGC target; 54-ms maximum injection time). Singly charged and unassigned precursors were excluded, and dynamic exclusion was set to 35 s.

Raw full-proteome data were processed using MaxQuant version 2.1.4.0^32^ against the human UniProt reference proteome containing one protein sequence per gene (20,650 entries; downloaded January 21, 2025). Default parameters were used with match between runs enabled, label-free quantification enabled^33^, and iBAQ calculation enabled.

### Differential proteomic and enrichment analysis

The MaxQuant proteinGroups output was filtered to remove potential contaminants and reverse hits. Zero LFQ intensities were treated as missing values. Each cell line was analyzed separately, with NTC as control and sg1-TRIM11 or sg2-TRIM11 as independent knockout conditions. LFQ intensities were processed using DEP (v. 1.28.0)^34^. Proteins with no more than one missing value in at least one condition were retained, followed by variance-stabilizing normalization^35^ and MinProb imputation (q = 0.01). Differential protein abundance was assessed separately for each TRIM11-targeting sgRNA using linear models with empirical Bayes moderation^36^, comparing each TRIM11-targeting condition with the non-targeting control (NTC). Proteins with an fdrtool q value (p.adj) < 0.05 were considered significant in each individual comparison. A grouped strip plot was used to summarize consensus differentially abundant proteins across the three cell lines. Consensus proteins were defined as proteins that were significant in both the sg1-TRIM11 versus NTC and sg2-TRIM11 versus NTC comparisons and showed the same direction of change. For visualization, the plotted log2 fold changes and fdrtool q values were calculated as the arithmetic means of the corresponding values from the two comparisons; these mean values were not used to determine statistical significance.

Gene Ontology over-representation analysis was performed using clusterProfiler (v. 4.14.6)^37^ and org.Hs.eg.db (v. 3.20.0) across all ontologies with Benjamini–Hochberg correction. Terms with adjusted P < 0.05 and at least four contributing proteins were used for network analysis. Enrichment networks were generated in Cytoscape 3.9.0^38^ with EnrichmentMap 3.5.0^39^. Edges represented overlap between term-associated protein sets and were retained at a Jaccard similarity of at least 0.25. Functionally related terms were grouped and annotated using AutoAnnotate 1.5.2^40^.

### Analysis of published RNA-sequencing and ChIP-sequencing data

For the RNA-sequencing comparison after EWSR1::ATF1 depletion, the published SUCCS1 count matrix from Ozenberger et al^10^ study was obtained from GEO series GSE248750. Counts were normalized using the trimmed mean of M values (TMM) method and converted to counts per million (CPM) using edgeR (v. 4.4.2)^41^.

For visualization of EWSR1–ATF1 and H3K27ac occupancy at the PCK1 locus, processed ChIP-sequencing bigWig files from Möller et al. were downloaded from GEO series GSE180187. Control and EWSR1–ATF1-depleted tracks were visualized at the PCK1 locus using a common genomic coordinate and display scale in Integrative Genomics Viewer (IGV), version 2.16.0^42^.

### PEPCK activity assay

PEPCK activity was measured using the PEPCK Activity Assay Kit (Abcam, #ab239714). Up to 2 × 10⁵ cells were lysed in 200 µl of ice-cold assay buffer for 10 min on ice. Lysates were centrifuged at 10,000 × g for 10 min at 4°C, and supernatants were collected. 40 µl of each sample was used per reaction. Samples were assayed in technical duplicates according to the manufacturer’s protocol. Absorbance at 570 nm was recorded kinetically at 37°C at 1-min intervals for 30 min using a SpectraMax microplate reader. PEPCK activity was normalized to total protein concentration measured by BCA assay.

### Targeted LC–MS/MS metabolite analysis

Frozen cell samples were extracted with 200 µl ice-cold acetonitrile/methanol/water (3:1:1, v/v/v) containing 15 mM ammonium acetate (pH 9), then mixed, sonicated, and subjected to two freeze–thaw cycles consisting of 1 min in liquid nitrogen and 5 min of thawing on ice. Extracts were centrifuged for 15 min at −9°C at the instrument’s maximum speed. Clarified supernatants were aliquoted for derivatization of TCA cycle intermediates.

For TCA cycle metabolites, 50 µl extract was derivatized with 3-nitrophenylhydrazine and 1-ethyl-3-(3-dimethylaminopropyl) carbodiimide in pyridine/ethanol for 20 min at 40°C, cooled, and quenched with 12% acetic acid. After centrifugation, derivatized metabolites were separated on an ACQUITY UPLC HSS T3 column (100 × 2.1 mm) at 40°C and detected on a QTRAP 6500+ system in negative-ion scheduled-MRM mode. Data were acquired and processed using SCIEX OS 4.2.0.1599. Quantified TCA cycle metabolites were normalized to the input cell number and reported as pmol per 10⁶ cells. Oxaloacetate was not quantified using the calibration procedure, its peak area was normalized to the input cell number.

### Stable-isotope tracing and GC–MS analysis

Cells were incubated for 24 h in glucose-free RPMI 1640 containing 15% dialyzed FBS (Gibco) and 11 mM [U-¹³C6] glucose (Sigma-Aldrich, #389374). Frozen cell samples were extracted on the culture plates with 1 ml 50% methanol containing ribitol as an internal standard. After 5 min on ice, cells were scraped and transferred to tubes, 500 µl chloroform was added, and samples were vortexed and incubated for 5 min on ice. Samples were centrifuged at 11,000 × g for 10 min; 350 µl of the upper polar phase was collected and dried under vacuum without heating.

Derivatized metabolites were analyzed using a Shimadzu TQ8040-NX GC–MS/MS system fitted with an Rxi-5Sil MS column (30 m × 0.25 mm × 0.25 µm; Restek). One microliter was injected at 280°C with a split ratio of 10. The oven program comprised 1 min at 65°C, a 6°C/min ramp to 220°C, a 20°C/min ramp to 330°C, and a 5-min bake-out at 330°C, with helium as carrier gas. The ion source and interface temperatures were 200°C and 280°C, respectively, and data were acquired in scan mode after a 5-min solvent delay. Natural-isotope-abundance-corrected peak areas of the selected labelled isotopologues were analyzed using DExSI version 1.11 [PMID: 29360933], normalized to total protein content determined by BCA assay. For results presentation, the normalized abundance of each selected labelled isotopologue in knockout samples was expressed relative to the corresponding nontargeted control samples.

### Extracellular flux analysis

Oxygen consumption rate (OCR) and extracellular acidification rate (ECAR) were measured using a Seahorse XFe96 Extracellular Flux Analyzer (Agilent Technologies). Cells were seeded at cell-line-specific densities 2 days before measurement to achieve approximately 80% confluence on the day of the experiment. The sensor cartridge was hydrated overnight in Seahorse XF Calibrant one day before the assay. On the assay day, cells were equilibrated in Seahorse XF RPMI medium supplemented with 1 mM sodium pyruvate, 2 mM L-glutamine, and 10 mM D-glucose 1 h at 37°C in a non-CO_2_ incubator.

For mitochondrial stress measurements, 1 µM oligomycin, FCCP (0.5 µM for CCS-SN1; 2 µM for other cell lines), and 0.5 µM rotenone plus 0.5 µM antimycin A were injected sequentially. For ECAR measurements, 1 µM oligomycin, FCCP (0.5 µM for CCS-SN1; 2 µM for other cell lines), 0.5 µM rotenone plus 0.5 µM antimycin A, and 75 mM 2-deoxy-D-glucose. For rescue experiments, cells were pretreated with 1 mM DM-αKG for 1 h before plate loading. After each assay, total protein per well was measured by BCA assay, and OCR and ECAR were normalized to protein content. Data were analyzed using Wave software, version 2.6.3 (Agilent Technologies).

ATP production rates were calculated from normalized OCR and Wave-processed glycoPER data according to the Agilent Seahorse XF Real-Time ATP Rate Assay Kit User Guide (Agilent Technologies, 2024). ATP-linked respiration was calculated as:

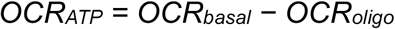

Mitochondrial ATP production was calculated using a phosphorylation-to-oxygen (P/O) ratio of 2.75:

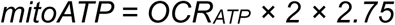

Glycolytic ATP production was calculated as glycoATP = glycoPER_basal_, and total ATP production as:

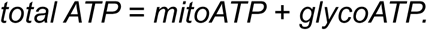

### Intracellular ATP measurement

Cells (1 × 10⁵ in 50 µl of medium per well) were seeded into white opaque 96-well plates and allowed to attach for 4 h. Intracellular ATP was measured using CellTiter- lo Luminescent Cell Viability Assay (Promega) according to the manufacturer’s instructions. An equal volume of CellTiter-Glo reagent was added, plates were shaken for 5 min and incubated for 10 min at room temperature, and luminescence was measured using a microplate reader. Signals from non-targeting control cells was used for normalization.

### Mitochondrial membrane potential and reactive oxygen species

Cells (2.5 × 10^5^ per well in 12-well plates) were stained at 37°C for 20 min in the dark with 2.5 µM DCF-DA (MedChemExpress, # HY-D0940), 5 µM MitoSOX Red (MedChemExpress, #HY-D1055), or 2.5 µg/ml JC-1 (MedChemExpress, #HY-K0601) according to the manufacturer’s instructions. Cells were washed twice with PBS and analyzed using a BD FACSCelesta flow cytometer.

### Annexin V/PI Apoptosis assay

Rescue cells were treated with 1 mM dimethyl α-ketoglutarate (DM-αKG) (Sigma-Aldrich, #349631) for 48 hours. Cells were then washed twice with PBS and resuspended in Annexin V Binding Buffer (BioLegend, #422201) at 2.5 × 10^6^ cells/ml. Aliquots of 100 µl were incubated with 5 µl fluorochrome-conjugated Annexin V and propidium iodide (BioLegend, #421301) for 15 min at room temperature in the dark. Following incubation, 400 µl of Annexin V Binding Buffer was added to each sample, and apoptosis was measured using a FACSCelesta SORP cell analyzer (BD Biosciences). Data were analyzed using (FlowJo v.X / BD FACSDiva software).

### Transmission electron microscopy

Cells grown on Aklar fluoropolymer film were fixed in 2% formaldehyde and 2% glutaraldehyde containing 1 mM CaCl2 and 1 mM MgCl2 in 100 mM sodium cacodylate buffer (pH 7.2), post-fixed in buffered 1% osmium tetroxide, stained in bloc with 1% uranyl acetate, dehydrated through graded ethanol, and embedded in epoxy resin. Ultrathin sections (nominal thickness, 60 nm) were contrast-stained with lead citrate and uranyl acetate and examined using a Zeiss EM 910 transmission electron microscope operated at 80 kV. Images were acquired with a slow-scan CCD camera (TRS, Moorenweis, Germany).

### Patient cohort and survival analysis

Tumor RNA-sequencing data from 29 matched CCS patient samples were integrated with the corresponding clinical data. Gene expression was represented as log2(TPM + 1). For Kaplan–Meier analyses, samples were divided into PCK1-high and PCK1-low groups using the cohort median PCK1 expression value (1.373 log2[TPM + 1]) as the cutoff. Progression-free survival analysis was restricted to 19 primary cases without metastasis at diagnosis, whereas overall survival analysis included all 29 samples. Survival curves were estimated using the Kaplan–Meier method and compared using the log-rank test; patients without an event were censored at their last recorded follow-up.

For progression-group comparisons, tumors were categorized as early progression, late progression, or metastatic at diagnosis according to the clinical annotation; Early and late progression were defined among primary cases without metastasis at diagnosis using the cohort median PFS time of approximately 900 days as the cutoff; tumors with PFS shorter than this cutoff were classified as early progression, whereas tumors with PFS longer than this cutoff were classified as late progression. The metastatic-at-diagnosis group included tumors from patients with metastatic disease present at initial diagnosis. Right-censored cases were excluded from this comparison. Overall group differences were assessed using the Kruskal–Wallis test and pairwise comparisons using two-sided Wilcoxon rank-sum tests. Associations between PCK1 and SNAI1 or MTSS1 expression were assessed using Spearman rank correlation. Analyses were performed using R (v. 4.4.3) with survival (v. 3.8-3), ggplot2 (v. 4.0.0), readxl (v. 1.4.5), and dplyr (v. 1.1.4).

### Pathway correlation analysis

Gene expression values were transformed as log2(TPM + 1). Gene sets from the Hallmark and Gene Ontology Biological Process, Molecular Function, and Cellular Component collections were obtained from the Molecular Signatures Database^43^ using msigdbr (v. 25.1.1). Gene sets containing 10–500 genes represented in the expression matrix were retained. Sample-wise pathway enrichment scores were calculated using GSVA (v. 2.0.7)^44^ with a Gaussian kernel. Spearman correlation analysis was performed between each pathway score and PCK1 expression across 29 CCS samples. P values were adjusted for multiple testing using the Benjamini– Hochberg method.

### Study Design and Patient Cohort

We conducted a retrospective cohort study of patients diagnosed with clear cell sarcoma who received treatment at sarcoma centres in the UK (Royal National Orthopaedic Hospital; London, Royal Orthopaedic Hospital (Arden Tissue Bank; Birmingham), Republic of Ireland, and Switzerland. Samples and data were also obtained from the Vivo Biobank (Newcastle UK); and the MCRC Biobank (Manchester, UK).

The study encompassed 29 tumor samples from 29 patients (age 21-56 years) collected between 1980-2026). The study was based on 28 samples from primary lesions, 2 from local recurrences, 1 from lymph node metastases and 1 from systemic metastases.

### Ethics Approval and Patient Consent

The Royal National Orthopaedic Hospital (RNOH) Biobank is approved by the National Research Ethics Committee of the Health Research Committee (reference: Integrated Research Application System (IRAS) project identifier: 272816). This study was approved by the UCL/UCLH Biobank for Studying Health and Disease – EC39.25 The Multi-Omic Diversity of Clear Cell Sarcoma. Written informed consent was obtained from all individual participants prior to inclusion in the study.

### Inclusion and Exclusion Criteria

All patients with a clinically confirmed diagnosis of clear cell sarcoma via standard clinical protocols (histology, IHC, EWSR1 FISH and/or RNA fusion panel, WGS) irrespective of primary site or disease stage were eligible for the study. All diagnoses were reviewed by a specialist sarcoma pathologist and diagnoses were revised based on results of histology and molecular tests. Patients were excluded based on lack of a detectable EWSR1 fusion or the patient declined biobank consent.

### Data Collection

Clinical and treatment data were extracted from NHS health records and through available biobank records which included at a minimum: 1) age at diagnosis, 2) date of diagnosis, 3) sex, 4) size of primary tumor, 5) site of primary tumor, 6) presence of metastasis at diagnosis, 6) type of synchronous metastasis, 7) type of fusion (ATF1, CREB1 or CREM) and any available surgical, radiotherapy or medical oncology regimens.

### End Points and Follow-Up

The primary survival endpoint was Overall Survival (OS), defined as the time from the time of disease diagnosis to death. Patients alive at the end of the study period were censored at the date of their last follow-up. Secondary endpoint was Progression Free Survival, defined as the time from disease initiation to the time of first disease progression or recurrence. Follow-up data were collected through NHS medical records up to May 2026.

## Declaration of generative AI and AI-assisted technologies

During the preparation of this manuscript, the authors used ChatGPT (OpenAI) and Claude (Anthropic) to improve the language, clarity, and readability of the manuscript. The authors reviewed and edited all AI-assisted text and take full responsibility for the content of the manuscript.

## Supplementary Tables

Table 1. Results of the BISON CRISPR/Cas9 knockout screen.

Table 2. Results of proteomics analysis comparing TRIM11 knockout and control in CCS cell lines.

Table 3. Results of the Pathways Network analysis.

Table 4. Clinical outcome data and correlation with PCK1 gene expression.

Table 5. Results of tumor transcriptomic programs correlated with PCK1 expression.

## Data and Code Availability

RNAseq data used in this study will are available via controlled access in the European Genome Archive under accession number (PLACEHOLDER, to be updated). Further details, including data access policy for the study, can be found there. No custom code was developed in the framework of this study.

## Acknowledgements

We sincerely thank Fenja Guyot and Abigail Ternynck for their excellent technical assistance. This work was supported by the Emmy Noether Program of the Deutsche Forschungsgemeinschaft (CH2301/1-1) to P.C., which also supported N.B., and by the HEROES-AYA consortium within the National Decade Against Cancer of the German Federal Ministry of Education and Research (01KD2207) to P.C. that supported K.K. and L.C. The clinical and tissue data collection and RNA sequencing was funded by the Edward Showler Foundation. A.F. and C.D receive funding support from the Edward Showler Foundation and Sarcoma UK respectively. N.P. holds a Cancer Research UK Career Establishment award (RCCCEA-Nov23/100003). The work in D.K. and S.G. are supported by the Stiftung Sarkomforschung (Sarcoma Research Foundation), Sarkom Tour (www.sarkomtour.de) and the Human sarcoma model initiative (www.husarc.org) grants to S.B.

## Authors contributions

W.C., M.Sl. and P.C. conceptualised the study. W.C., K.B., K.K., M.Sl. and P.C. designed the experiments.. W.C., K.B., K.K., L.M., J.P., N.B., L.C. and A.R. performed the experiments. W.C., K.B., K.K., L.M., J.P., A.F., N.B. and P.C. analysed and interpreted the data. D.K., N.B., A.D., K.S.L. and F.Y.-R. advised on key experimental procedures. M.Sch., D.F. and S.A. supported proteomics data acquisition and analysis. M.Sa. and M.B. supported metabolomics data acquisition and analysis. K.R. and M.N. supported transmission electron microscopy. A.F., C.D., D.H., S.S. and N.P. collected and curated biospecimens, tumor RNA sequencing and clinical data. D.K, S.G. and S.B. contributed model systems through HUSARC Sarcoma Model Initiative. C.A.O, C.S. and S.F. provided access to key laboratory equipment and infrastructure. G.M., A.Y.R., C.A.O, K.J., N.P. and S.B. contributed expert guidance. P.C. supervised the study and acquired funding, with supporting contributions from N.P. and S.B. W.C. and P.C. wrote the manuscript with input from all coauthors. All authors read and approved the final manuscript.

