## Supplementary Figure S1 for "EWSR1::ATF1 hyperactivation via TRIM11 depletion triggers lethal metabolic rewiring in clear cell sarcoma"

### Supplementary Figure 1

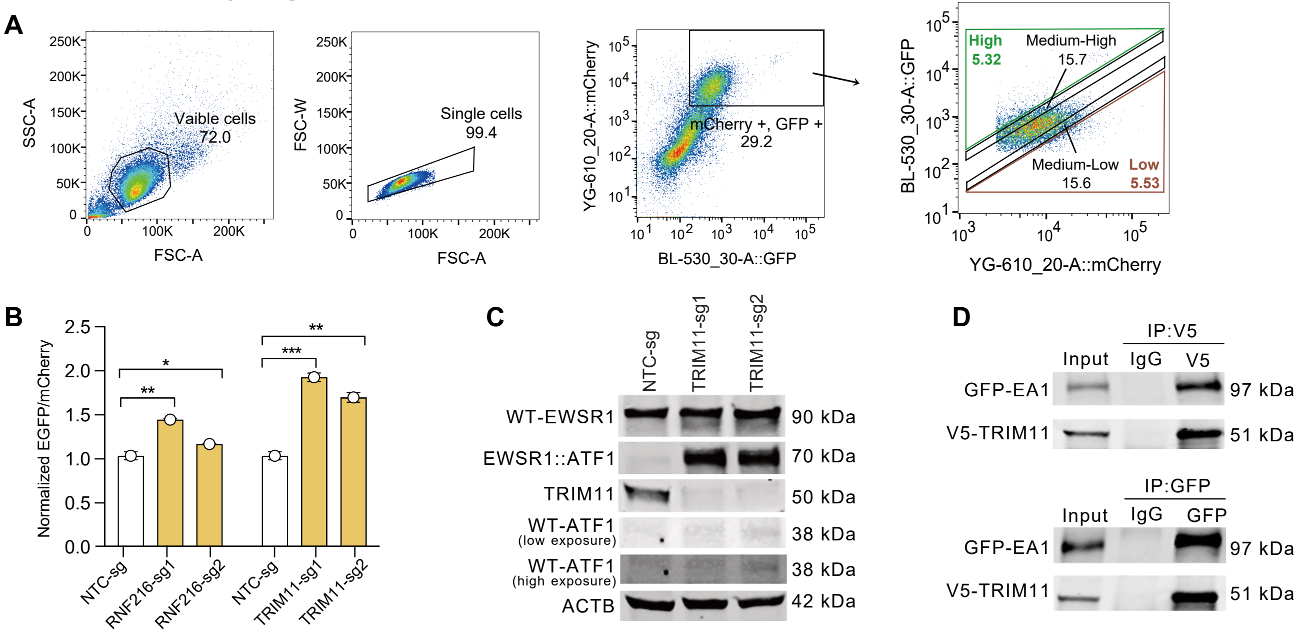


**Figure S1. A ubiquitin–proteasome system (UPS)-focused CRISPR screen identifies TRIM11 as a regulator of EWSR1::ATF1 fusion stability.**

(A) Flow cytometric sorting strategy for the Bison CRISPR/Cas9 screen. To enrich for HEK293T Cas9 cells expressing the EWSR1::ATF1 reporter, viable singlets were first selected, followed by gating of GFP- and mCherry-double-positive cells. Double-positive cells were then separated according to their GFP/mCherry fluorescence ratio into GFP-high (upper approximately 5%), GFP-medium-high (adjacent approximately 15%), GFP-medium-low (adjacent approximately 15%), and GFP-low (lower approximately 5%) fractions. The GFP-high and GFP-low fractions were used for single-guide RNA (sgRNA) enrichment analysis. CRISPR, clustered regularly interspaced short palindromic repeats.

(B) Flow cytometric analysis of fusion enrichment upon RNF216 knockout (RKO) and TRIM11 knockout (TKO) in HEK293T with EWSR1::ATF1 pArtichoke. P values were calculated using ordinary one-way ANOVA followed by Dunnett's multiple comparisons test against the non-targeting control (NTC) within each knockout condition *P < 0.05; **P < 0.01; ***P < 0.001. Data are representative of two biological replicates.

(C) Western blot analysis of wild-type EWSR1 (WT-EWSR1) and wild-type ATF1 (WT-ATF1) upon TKO compared with NTC in CCS210.

(D) Co-immunoprecipitation analysis of the interaction between GFP-tagged EWSR1::ATF1 (GFP-EA1) and V5-tagged TRIM11 in CCS210 cells. Cell lysates were subjected to immunoprecipitation with an anti-V5 antibody or control IgG, followed by immunoblotting for GFP, V5, and beta-actin. Whole cell lysate is used as input.
