## Supplementary Figure S2 for "EWSR1::ATF1 hyperactivation via TRIM11 depletion triggers lethal metabolic rewiring in clear cell sarcoma"

### Supplementary Figure 2

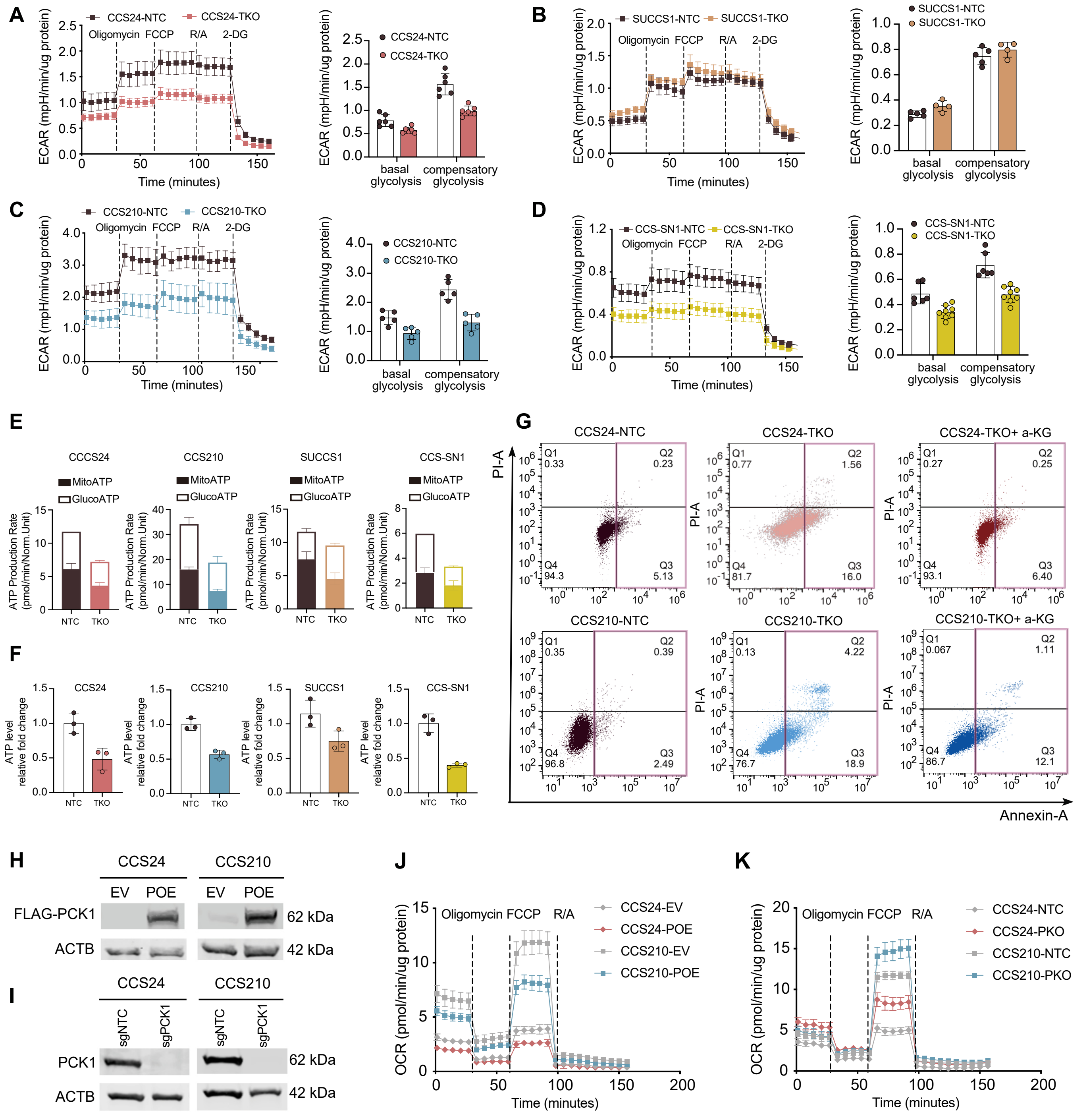


**Figure S2: TRIM11 knockout reduced cellular respiration with mitochondrial damage**

(A-D) Seahorse analysis of the extracellular acidification rate (ECAR) in all four CCS cell lines with NTC and TRIM11 knockout (TKO). Vertical dashed lines indicate the sequential addition of the indicated compounds. FCCP, carbonyl cyanide-p-trifluoromethoxyphenylhydrazone; R/A, rotenone/antimycin A; 2-DG, 2-deoxy-D-glucose.

(E) Seahorse analysis of ATP production rate of NTC and TKO in four CCS cell lines.

(F) Intracellular ATP abundance was measured using the CellTiter-Glo (CTG) assay in NTC and TKO cells of four CCS cell lines (n = 3 biological replicates).

(G) Flow cytometric analysis of cell viability using Annexin V/PI staining in CCS24 and CCS210 cells with NTC, TKO and TKO rescued by dimethyl-α-ketoglutarat (DM-α-KG). Colored lines highlight the apoptotic cell populations. The percentages of early apoptotic (Q3) and late apoptotic (Q2) cells are indicated.

(H) Western blot analysis of PCK1 expression in CCS24 and CCS210 cells transduced with an empty vector (EV) or a construct encoding FLAG-tagged PCK1 (POE).

(I) Western blot analysis of PCK1 expression in CCS24 and CCS210 cells transduced with a non-targeting control sgRNA (sgNTC) or a PCK1-targeting sgRNA (sgPCK1).

(J-K) Seahorse analysis of the oxygen consumption rate (OCR) in CCS24 and CCS210 with control and POE (C), with control and PKO (D). Vertical dashed lines indicate the sequential addition of the indicated compounds. FCCP, carbonyl cyanide-p-trifluoromethoxyphenylhydrazone; R/A, rotenone/antimycin A.
